# Alterations in human gut microbiome composition and metabolism after exposure to glyphosate and Roundup and/or a spore-based formulation using the SHIME® technology

**DOI:** 10.1101/2021.12.16.472928

**Authors:** Robin Mesnage, Marta Calatayud, Cindy Duysburgh, Massimo Marzorati, Michael N Antoniou

**Affiliations:** Gene Expression and Therapy Group, King’s College London, Faculty of Life Sciences & Medicine, Department of Medical and Molecular Genetics, Guy’s Hospital, London, SE1 9RT, UK; ProDigest BV, Technologiepark 82, 9052 Ghent, Belgium; Center For Microbial Ecology and Technology, Faculty of Bioscience Engineering, Department of Biotechnology, Coupure Links 653, 9052 Ghent, Belgium

## Abstract

Despite extensive research into the toxicology of the herbicide glyphosate, there are still major unknowns regarding its effects on the human gut microbiome. As a step in addressing this knowledge gap, we describe for the first time the effects of glyphosate and a Roundup glyphosate-based herbicide on infant gut microbiota using SHIME technology, which mimics the entire gastrointestinal tract. SHIME® microbiota culture was undertaken in the presence of a concentration of 100 mg/L (corresponding to a dose of 1.6 mg/kg/day) glyphosate and the same glyphosate equivalent concentration of Roundup, which is in the range of the US chronic reference dose, and subjected to molecular profiling techniques to assess outcomes. Roundup and to a lesser extent glyphosate caused an increase in fermentation activity, resulting in acidification of the microbial environment. This was also reflected by an increase in lactate and acetate production concomitant to a decrease in the levels of propionate, valerate, caproate and butyrate. Ammonium production reflecting proteolytic activities was increased by Roundup exposure. Global metabolomics revealed large scale disturbances in metabolite profiles, including an increased abundance of long chain polyunsaturated fatty acids (n3 and n6). Although changes in bacterial composition measured by qPCR and 16S rRNA sequencing were less clear, our results suggested that lactobacilli had their growth stimulated as a result of microenvironment acidification. Co-treatment with the spore-based probiotic formulation MegaSporeBiotic reverted some of the changes in short-chain fatty acid levels. Altogether, our results suggest that glyphosate can exert effects on human gut microbiota at permitted regulatory levels of exposure, highlighting the need for epidemiological studies aimed at evaluating the effects of glyphosate herbicides on human gut microbiome function.

## INTRODUCTION

Glyphosate is globally the most widely used broad-spectrum herbicide and crop desiccant (Maggi et al. 2019) being an active ingredient in many herbicide formulations, including Roundup. In addition to glyphosate salts, formulations of glyphosate-based herbicides (GBHs) contain a number of co-formulants such as surfactants, which vary in nature and concentration between different commercial products (Mesnage et al. 2019). These surfactants are added as wetting agents to maximize coverage and aid penetration of the herbicide through the plant cell walls. Glyphosate herbicides are generally used to kill weeds, especially annual broadleaf weeds and grasses that compete with crops. Although GBHs have been approved by regulatory bodies worldwide, concerns about their effects on humans and the environment persist (Robinson et al. 2020).

Human exposure to glyphosate has been described mainly through the oral route, direct exposure in occupational settings or environmental exposure through residues in food (Faniband et al. 2021a). From ingested glyphosate, oral absorption has been estimated in animal studies at approximately 20% (Anadón et al. 2009), with main excretion via faeces. Recently, a total dose recovery of glyphosate in urine has been described as low as 1-6% in human young adults, suggesting that glyphosate can reach the colonic environment and potentially affect the gut microbial ecosystem (Faniband et al. 2021b).

Glyphosate principally acts as a herbicide in plants by inhibiting the activity of 5-enolpyruvylshikimate-3-phosphate synthase (EPSPS) of the shikimate pathway (Boocock and Coggins 1983), causing a shortage in aromatic amino acid biosynthesis. The shikimate pathway is also found in microorganisms including those inhabiting the human gut microbiome (Mesnage and Antoniou 2020) As a consequence, concerns have been raised about the ability of glyphosate and GBHs to have adverse effects through its interactions with the community of bacteria residing in the digestive tract. The number of animal studies investigating the effects of glyphosate and GBHs on gut microbiota is growing (Tsiaoussis et al. 2019). We have recently described how glyphosate and a European Union Roundup GBH (MON 52276) can indeed inhibit EPSPS of the shikimate pathway in the gut microbiome of rats (Mesnage et al. 2021b). However, the gut microbiota of laboratory rodents or wild animals is substantially different from the human gut microbiota and knowledge gaps remain as to whether glyphosate also affects human gut microbiota.

Other chemical residues and environmental pollutants have been described to have a significant impact on human gut microbiota, likely affecting the host’s health status through modulation of metabolic, endocrine, and neurodevelopmental paths linked to the gut microbiome (Calatayud Arroyo et al. 2021). Infants are especially sensitive to xenobiotic exposures, and cumulative effects might be aggravated due to longer exposure times, putting them at a higher level risk compared to adults.

The first aim of the present study was to assess the impact of glyphosate and a GBH on the activity and composition of microbiota obtained from a 3-year-old human donor. We used SHIME® technology, which mimics the entire gastrointestinal tract by employing a series of reactors representing the different stages of food uptake and digestion (Venema and van den Abbeele 2013). This model system allows the culture of complex gut microbiota over a longer period under representative conditions of different intestinal regions. Therefore, SHIME® does not only allow the gathering of detailed information concerning fermentation profile, but importantly also about the localization of the intestinal fermentation activity. A testament to the usefulness of SHIME® technology in studying effects of chemical pollutants on the human gut microbiota, is its successful application in the study of chronic exposure to chlorpyrifos, which showed that this pesticide causes dysbiosis (Joly et al. 2013).

Probiotics have been traditionally used to modulate gut microbial communities and provide benefit to the host via different mechanisms (Astolfi et al. 2019). In addition to metabolite modulation, microbe-microbe interplay or host-microbe interactions, some probiotic strains have also been proven to reduce toxicity of xenobiotics by modulating oxidative stress, or xenobiotics adsorption capacity, with a focus on inorganic compounds of Pb, Cd, As and Hg (de Matuoka e Chiocchetti et al. 2020; Majlesi et al. 2017). However, no study has been performed to date to determine if the effects of glyphosate and GBHs on the gut microbiome can also be mitigated through the use of pre- and probiotics. Thus, a second aim of this study was to assess the efficacy of a spore-based formulation, MegaSporeBiotic (Microbiome Labs; https://microbiomelabs.com/home/), in remediating the effects of glyphosate on gut microbiota. Several studies have explored potential health benefits of *Bacillus* strains such as stimulation of innate immunity (Ciprandi et al. 1986; Ciprandi et al. 2005). In the case of *B. subtilis* var. three strains of *B. subtilis* have been registered with the FDA with self-affirmed GRAS status in the USA in 2012. Recently, an *in vitro* study using MegaSporeBiotic supplementation for 3 weeks showed a significant capacity to modulate microbial activity and structure, increasing bacterial diversity, propionate, and *Akkermansia muciniphila*, bifidobacteria, and Firmicute levels (Marzorati et al. 2021).

The human intestinal tract harbours a large and complex community of microbes, which is involved in maintaining human health by preventing colonization by pathogens, producing nutrients and signalling molecules that module the function of numerous physiological processes (Ducarmon et al. 2019). Microorganisms are not randomly distributed throughout the intestine and those adhering to the gut wall play an important role as a ‘barrier’ against pathogens, instructing mucosal immune responses and occupying a niche at the expense of potentially harmful colonizers. In order to also model both the luminal and mucosal microbial community, we used an adaptation of M-SHIME®, which takes colonization of the mucus layer into account (Van den Abbeele et al. 2013b).

Exposure to the different co-formulants included in commercial GBH, or added to these formulations as adjuvants to the herbicide spray tank mix, are a major source of occupational health hazards (Mesnage et al. 2019). For instance, Roundup (MON 2139) is used as a 2% solution (7.2 g/L of glyphosate, 3.6% surfactant MON 0818, which is mostly made of polyoxyethylene tallow amine) on most perennials weeds. Recent studies have suggested that GBH formulations can cause changes in gut microbiota composition, which are not detected with glyphosate alone. In a recent study combining shotgun metagenomics and metabolomics to evaluate effects of glyphosate and the EU representative GBH formulation Roundup MON 52276 in rats, *Shinella zoogleoides* was found to be increased by exposure to MON 52276 but not glyphosate (Mesnage et al. 2021b). This was linked to changes in alkaloid levels in the gut, as suggested by the large decrease in solanidine levels, which was only detected in the MON 52276 treated group. Therefore, a third aim of this study was to evaluate if glyphosate formulated products have different effects in comparison to glyphosate alone. For this purpose, the model was operated as a TWINSHIME® in order to test both glyphosate and a formulated GBH namely Roundup MON 76207.

## MATERIAL AND METHODS

### Chemicals

The test products were glyphosate certified reference material, 1000 μg/mL in H_2_O (EPA 547 Glyphosate Solution, Merck KGaA, Darmstadt, Germany) and the glyphosate containing herbicide formulation MON 76207 (Roundup PROMAX on the US market, EPA Registration No. 524-579). The daily dose of glyphosate was selected during a pre-screening experiment made to evaluate changes in microbial activity and *Enterobacteriaceae* levels, and set at a concentration of 100 mg/L. Roundup was tested at a similar glyphosate level, taking into account that Roundup consisted for 48.7% of glyphosate. During the co-treatment period, the spore-based probiotic, MegaSporeBiotic (Microbiome Labs Physicians Exclusive, LLC., FL, USA), was additionally administered at a concentration of 340 mg containing 4 billion CFU, corresponding to a human ingestion of 2 capsules/day. MegaSporeBiotic contains 5 different Bacillus strains; i.e., *Bacillus indicus HU36™*, *Bacillus clausii (SC-109), Bacillus subtilis HU58™*, *Bacillus licheniformis* (SL-307) and *Bacillus coagulans (SC-208)*.

### Fecal sample

An infant donor was selected based on the following inclusion criteria: healthy, age between 12-36 months, no antibiotics or any other drug intake at least during the previous six months, normal weight, no constipation, hospital-born, no known diseases at least during the previous six months, exclusive breastfeeding for at least three months. The fecal sample was obtained in a plastic container with an “Oxoid™ AnaeroGen™” bag to limit the sample’s exposure to oxygen, immediately transported to the lab and used to inoculate the M-SHIME system. Briefly, a mixture of 1:10 (w/v) of fecal sample and anaerobic phosphate buffer (K_2_HPO_4_ 8.8 g/L; KH_2_PO_4_ 6.8 g/L; sodium thioglycolate 0.1 g/L; sodium dithionite 0.015 g/L; N2-flushed for 15 min) was homogenized for 10 min (BagMixer 400, Interscience, Louvain-La-Neuve, Belgium). After centrifugation (500g, 2 min) (Centrifuge 5417C, Eppendorf, VWR, Belgium) to remove large particles, the fecal sample was used to inoculate different M-SHIME reactors.

### Short-term colonic incubation

Short-term colonic incubations were performed to determine the concentration of glyphosate to be administered in the long-term SHIME experiment. The short-term screening assay tested a concentration range of glyphosate (0, 0.5, 1, 3, 10, 100 and 1000 mg/L) on a simulated proximal colonic environment (pH 6.2-6.4) with a representative bacterial inoculum. This bacterial inoculum was obtained from the long-term M-SHIME reactor during the stabilization period and simulated the colonic microbiota of a 3 year old infant.

Briefly, short-term colonic incubations were performed by inoculating a 10% (v/v) of a homogeneous mixture of the ascending, descending and transverse colon M-SHIME luminal fluid (stabilization phase) into colonic media (K_2_HPO_4_ 5.7 g/L; KH_2_PO_4_ 17.9 g/L; NaHCO_3_; 2.2 g/L; yeast extract 2.2 g/L; peptone 2.2 g/L; mucin 1.1 g/L; cysteine 0.5 g/L; Tween80 2.2 mL/L). Details on the M-SHIME inoculum are described in the next section. Incubations containing different glyphosate concentrations were performed for 48h at 37°C, under shaking (90 rpm) and anaerobic conditions. Short-term effects of different glyphosate concentrations on microbial activity markers (gas production, acid/base consumption, lactate, ammonia, short chain fatty acids, and branched fatty acids) and qPCR measurement of *Enterobacteriaceae*, was assessed at time points 0, 6, 24 and 48h. Since the production of microbial metabolites in the colon reactors alters the pH, the pH is controlled through the addition of acid or base (i.e. base consumption).

### Long-term colonic incubation

The reactor setup of the M-SHIME®, representing the gastrointestinal tract of the infant human was conducted as first described by Molly and colleagues (Molly et al. 1993), with modifications as defined by Van den Abbeele et al.(Van den Abbeele et al. 2013a; Van den Abbeele et al. 2012; Van den Abbeele et al. 2021).

The M-SHIME included both the luminal and mucosal microbiota, and consisted of three colonic vessels simulating the ascending, transverse and descending colon. Anaerobic conditions were achieved by 15 min flushing of all reactors with N2, 15 min twice per day. Double jacketed reactors connected to a recirculating warm water bath kept the M-SHIME units at 37°C and continuous stirring was maintained throughout the duration of the experiment. All reactors incorporated a mucosal environment in the luminal suspension. The mucosal environment consisted of 80 mucin agar-covered microcosms (AnoxKaldnes K1 carrier; AnoxKaldnes AB, Lund, Sweden) per vessel, prepared and submerged in defined colonic medium as previously described (Calatayud et al. 2021).

Each M-SHIME reactor was inoculated with 20% (v/w) infant fecal inoculum. During the first 16 h, reactors were operated in batch mode to allow for initial stabilization of the system and colonization of the mucus microenvironment. Subsequently, the stabilization period (2 weeks, day −14 to day −1; Figure S1) started in semi-continuous mode. Each experimental M-SHIME unit consisted of a first reactor that simulated over time the stomach and small intestine and that operated according to a fill-and-draw principle, with peristaltic pumps adding a defined amount of nutritional colonic medium to the stomach (gastric reactor = 140 mL; pH 3; 30 min), followed by addition of 60 mL of simulated pancreatic and bile juice (NaHCO_3_ 2.6 g/L, Oxgall 4.8 g/L and pancreatin 1.9 g/L) (small intestinal reactor = 200 mL; pH 6.5; 105 min). After this time, the intestinal suspension was pumped to the ascending colon vessel (AC reactor = 500 mL, pH 5.7-5.9; 250 min), and sequentially to the transverse (TC reactor = 800 mL, pH 6.2-6.4; 260 min) and descending colon (DC reactor = 600 mL; pH 6.6-6.9; 280 min) vessels. The pH was automatically adjusted by continuous measurement (Senseline pH meter F410; ProSense, Oosterhout, The Netherlands) coupled to a pH controller (Consort R301) and automatic pump (Master Flex 109 pump drive; Cole-Parmer Instrument Company, LLC), with dosing of either HCl (0.5 M) or NaOH (0.5 M) (Carl Roth, Belgium), as required.

After the stabilization period, a control period (C) of two weeks (d0 to d14) was used as a baseline for microbial community composition and activity (Figure 1). During the control period, samples were obtained three times per week from the luminal environment to evaluate microbial activity (measurement of lactate, ammonia, short chain fatty acids and branched fatty acids), and once per week for community composition employing qPCR for detection of Firmicutes and Bacteroidetes phyla, *Lactobacillus* spp., *Bifidobacterium* spp., *Akkermansia muciniphila*, and *Faecalibacterium prausnitzii*, and 16S rRNA Illumina sequencing. Community composition of the mucosal environment was also analysed in samples obtained once per week from the mucus beads.

**Figure 1.**
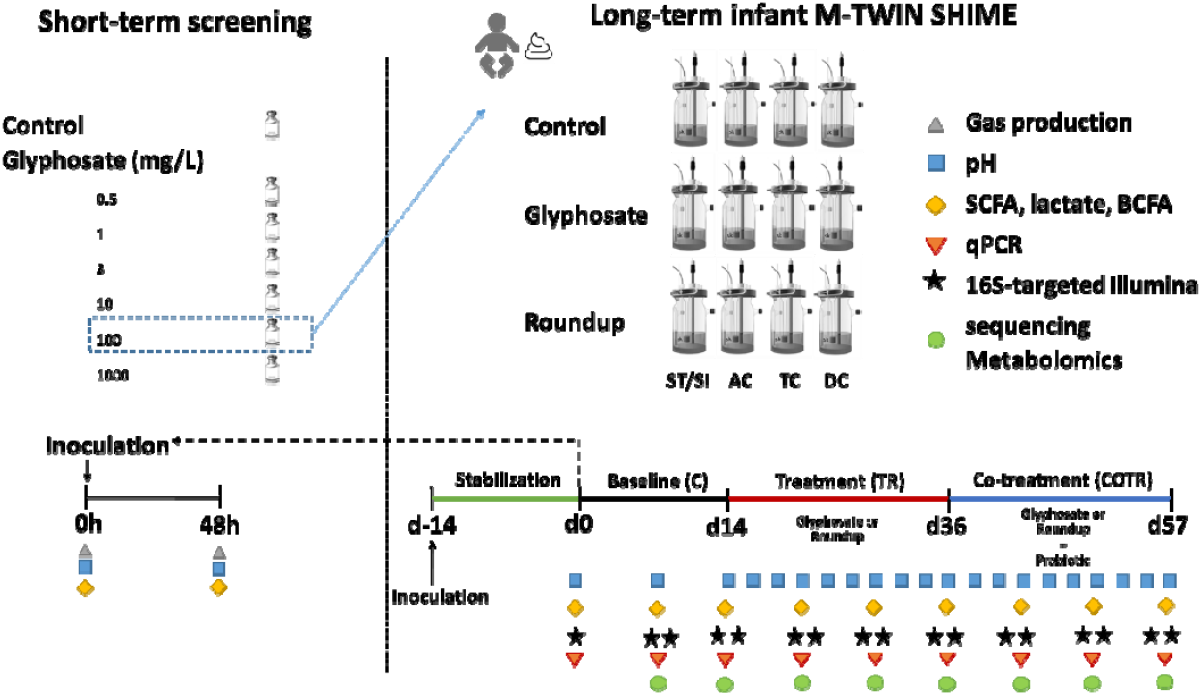
Experimental design. TWINSHIME® was used in order to evaluate the impact of glyphosate and a Roundup glyphosate-based herbicide formulation on the microbiota of a healthy human 3-year-old donor. The effect of adding MegaSporeBiotic spore-based probiotic. Culture period was for 10 weeks in chambers simulating the ascending (AC), transverse (TC) and descending colon (DC). The index highlights the molecular measurements undertaken to assess the effects of the different treatments at various sampling timepoints (lower right panel).

Subsequently, during the treatment period (TR), a concentration of 100 mg/L glyphosate as pure standard or Roundup herbicide formulation at the same glyphosate equivalent concentration was added to the colonic nutritional media, and administered to the stomach reactor three times per day over three weeks, corresponding to a dose of glyphosate of 14 mg to be distributed over the SHIME unit (AC, TC and DC, total volume = 1.9L). Due to the dynamic nature of the system following a fill and draw principle, a steady-state equilibrium was expected after the first 72h of the assay. After TR period, supplementation with the MegaSporeBiotic spore-based probiotic (4 billion CFUs) was administered on top of the nutritional media and glyphosate (COTR) for three more weeks (Figure 1).

During TR and COTR periods, samples were obtained three times per week for assessing microbial metabolic activity and once per week for microbial community composition, as previously described (Figure 1). Additionally, luminal samples for metabolomics were obtained once per week during C, TR and COTR periods.

### Microbial community activity

Metabolic activity of the gut microbial communities was evaluated by quantifying general markers of fermentation (pH and gas production), lactate, ammonia, short chain fatty acid and branched fatty acid production during the short-term experiment at 0, 6, 24 and 48h. During the long-term M-TWIN SHIME assay, the same markers were quantified three times per week, with the exception of gas production.

The pH was recorded using an automatic probe (Senseline F410; ProSense, Oosterhout, The Netherlands). Gas production was estimated by measuring bioreactor pressure with a hand-held pressure indicator (CPH6200; Wika, Echt, The Netherlands). Lactate quantification was performed using a commercial enzyme-based assay kit (R-Biopharm, Darmstadt, Germany) according to the manufacturer’s instructions. Short-chain fatty acids (SCFAs) and branched fatty acids (b-SCFAs) measurements were conducted as previously described (De Weirdt et al. 2010). Ammonia was quantified by the Kjeldahl nitrogen determination method again as previously described (Chaikham et al. 2012).

### DNA extraction

Total genome DNA from pelleted bacterial cells obtained from a 1□mL sample or 0.25 □g mucin agar collected from the mucus beads, was extracted as previously described (Boon et al. 2003) with modifications as described by Dayshburg and colleagues (Boon et al. 2003; Duysburgh et al. 2019). DNA concentration and purity was monitored by electrophoresis on 2% (w/v) agarose gels and spectrophotometrically by determination of A260/A280 ratios (Synergy HT Microplate Reader, BioTek, USA). Amplicon based metagenomics was performed using the NEBNext® Ultra™ IIDNA Library Prep Kit (Cat No. E7645).

### Microbial community composition via qPCR

Specific taxa (Firmicutes and Bacteroidetes phylum, *Enterobacteriaceae* family, *Bifidobacterium* spp., *Lactobacillus* spp., *Akkermansia muciniphila* and *Faecalibacterium prausnitzii*) were analysed by qPCR using the primers and conditions described in Supplementary Table S1 and S2. The qPCR was performed using a QuantStudio 5 Real-Time PCR system (Applied Biosystems, Foster City, CA, USA). Each sample was run in technical triplicate and results are reported in log bsed format (16S rRNA gene copies/mL).

### 16S rRNA gene amplicon sequencing

A total of 200ng DNA was used for PCR amplification with primer sets targeting different 16S rRNA gene hypervariable regions. Microbial community composition was assessed at defined time points by next-generation 16S rRNA gene amplicon sequencing of the V3–V4 region under contract with Novogene (UK) Company Limited (Cambridge, UK). Primers targeting the V3-V4 hypervariable region of bacterial 16S rRNA genes were as follows: 341F (5′-CCTAYGGGRBGCASCAG-3′) and 806R (5′-GGACTACNNGGGTATCTAAT-3′), where F is forward and R is reverse. Each primer set was ligated with a unique barcode sequence. The PCR product was then selected for proper size and purified for library preparation. The same amount of PCR product from each sample was pooled, end polished, A-tailed, and ligated with adapters. After purification, the library was analyzed for size distribution, quantified using real-time PCR, and sequenced on a NovaSeq 6000 SP flowcell with PE250 platform.

### Metabolomics

All samples were maintained at −80°C until processed. Samples were prepared using the automated MicroLab STAR® system from Hamilton Company. To remove protein, dissociate small molecules bound to protein or trapped in the precipitated protein matrix, and to recover chemically diverse metabolites, proteins were precipitated with methanol under vigorous shaking for 2 min (Glen Mills GenoGrinder 2000) followed by centrifugation. The resulting extract was divided into five fractions: two for analysis by two separate reverse phase (RP)/UPLC-MS/MS methods with positive ion mode electrospray ionization (ESI), one for analysis by RP/UPLC-MS/MS with negative ion mode ESI, one for analysis by HILIC/UPLC-MS/MS with negative ion mode ESI, and one sample was reserved for backup. Samples were placed briefly on a TurboVap® (Zymark) to remove the organic solvent. The sample extracts were stored overnight under nitrogen before preparation for analysis. The details of the solvents and chromatography used are already described (Ford et al. 2020).

All methods utilized a Waters ACQUITY ultra-performance liquid chromatography (UPLC) and a Thermo Scientific Q-Exactive high resolution/accurate mass spectrometer interfaced with a heated electrospray ionization (HESI-II) source and Orbitrap mass analyzer operated at 35,000 mass resolution. The sample extract was dried then reconstituted in solvents compatible to each of the four methods. Each reconstitution solvent contained a series of standards at fixed concentrations to ensure injection and chromatographic consistency. One aliquot was analyzed using acidic positive ion conditions, chromatographically optimized for more hydrophilic compounds. In this method, the extract was gradient eluted from a C18 column (Waters UPLC BEH C18-2.1×100 mm, 1.7 μm) using water and methanol, containing 0.05% perfluoropentanoic acid (PFPA) and 0.1% formic acid (FA). Another aliquot was also analyzed using acidic positive ion conditions, however it was chromatographically optimized for more hydrophobic compounds. In this method, the extract was gradient eluted from the same aforementioned C18 column using methanol, acetonitrile, water, 0.05% PFPA and 0.01% FA and was operated at an overall higher organic content. Another aliquot was analyzed using basic negative ion optimized conditions using a separate dedicated C18 column. The basic extracts were gradient eluted from the column using methanol and water, however with 6.5mM ammonium bicarbonate at pH 8. The fourth aliquot was analyzed via negative ionization following elution from a HILIC column (Waters UPLC BEH Amide 2.1×150 mm, 1.7 μm) using a gradient consisting of water and acetonitrile with 10mM ammonium formate, pH 10.8. The MS analysis alternated between MS and data-dependent MSn scans using dynamic exclusion. The scan range varied slightly between methods but covered 70-1000 m/z. Raw data files are archived and extracted as described below.

Raw data was extracted, peak-identified and QC processed using Metabolon’s hardware and software as already described (DeHaven et al. 2010). Several controls were analyzed in concert with the experimental samples and were used to calculate instrument variability (5%) and overall process variability (7%). Peak area values allowed the determination of relative quantification among samples (Evans et al. 2009). The present dataset comprises a total of 842 biochemicals, 721 compounds of known identity (named biochemicals) and 121 compounds of unknown structural identity (unnamed biochemicals).

### Bioinformatics

Data processing was performed on the Rosalind High Performance Compute (HPC) cluster (a BRC/King’s College London computing facility to quickly perform large-scale calculations with a virtual machine cloud infrastructure) using 4 threads and a maximum RAM of 32 GB. The DADA2 algorithm (version 1.6) was first used to correct for sequencing errors and derive amplicon sequence variants (ASV) using the R version 3.4.1. ASV are a higher-resolution version of the operational taxonomic unit (OTU). While OTUs are clusters of sequences with 97% similarity, ASVs are unique sequences (100% similarity). The use of ASV is being advocated to replace OTU in order to improve reproducibility between studies because a threshold of 97% does not allow a reproducible classification of closely related bacteria (Callahan et al. 2017).

The taxonomy was assigned using the SiLVA ribosomal RNA gene database v132. The count table, the metadata, and the sequence taxonomies were ultimately combined into a single R object, which was analysed using the phyloseq package (McMurdie and Holmes 2013). The alpha diversity, representing the diversity of the total number of species (the richness R) within the samples, was measured using the Shannon’s diversity index (Morgan and Huttenhower 2012). Another important diversity measure is the beta diversity, representing the diversity of the total number of species between the samples (Morgan and Huttenhower 2012). We used the Bray-Curtis dissimilarity index, which was calculated using the Vegan R package (version 2.5.2). The analysis of 16S rRNA gene composition was conducted with a linear-mixed model with MaAsLin (Microbiome Multivariable Association with Linear Models) 2.0 (package version 0.99.12) (Mallick et al. 2021). The taxonomic composition was transformed using an arcsine square root transformation. The Benjamini–Hochberg method was used to control the False Discovery Rate (FDR) of the MaAsLin analysis.

We further analysed the 16S rRNA gene sequencing dataset and the metabolome dataset with an orthogonal partial least squares discriminant analysis (OPLS-DA), which are robust methods to analyse large datasets in particular when the number of variables is larger than the number of samples. OPLS-DA can be used to distinguish the variability corresponding to the experimental perturbation from the portion of the data that is orthogonal; that is, independent from the experimental perturbation. The R package ropls version 1.20.0 was used with a nonlinear iterative partial least squares algorithm (NIPALS) (Thévenot et al. 2015). Prior to analysis, experimental variables were centred and unit-variance scaled. Since PLS-DA methods are prone to overfitting, we assessed the significance of our classification using permutation tests (permuted 1,000 times). The variables of importance (VIP) were extracted from the models to determine as to what are the effects of the treatments.

In addition, Calypso version 8.4 (Zakrzewski et al. 2017) and MicrobiomeAnalyst (update 10/18/2021) online software (Dhariwal et al. 2017) were used to perform discriminant analysis of principal components (DAPC), linear discriminant analysis effect size (LEfSe), heat tree (Foster et al. 2017) and a zero inflated Gaussian fit mix model from 16S rRNA gene sequence data normalized by Cumulative-sum scaling (Paulson et al. 2013) (CSS) and log2 transformation to account for the non-normal distribution of taxonomic counts data.

## RESULTS

### Pre-screening experiment to determine glyphosate dosage

A first experiment was performed to test the short-term effect of 6 different concentrations of glyphosate (0.5, 1, 3, 10, 100 and 1000 mg/L) on microbial activity and *Enterobacteriaceae* levels. The European Food Safety Agency (EFSA) has established an acute reference dose (ARfD) and acceptable daily intake (ADI) for glyphosate of 0.5 mg/kg body weight/day (EFSA 2015). Assuming 15-20 kg of an infant of 3 years old, this would correspond to 7.5-10 mg/day, proving the doses used in this study are regulatory relevant.

Monitoring the pH during a colonic incubation provides an overall indication of the microbial fermentative activity (Supplementary Table 3). Limited effects of glyphosate were observed although glyphosate at a concentration of 1000 mg/L lowered the acidity of the medium, which could indicate a modulatory effect on microbial activity resulting in altered metabolite (i.e., lactate, acetate) production. Total gas production was also measured to reflect the rate of substrate fermentation (Supplementary Table 3). Glyphosate tended to slightly decrease the amount of gas production, except at a concentration of 1000 mg/L for which an increased gas production was observed. SCFA production was affected by the highest concentration of glyphosate. A lower production of acetate, propionate and branched SCFAs and higher levels of butyrate were observed compared to the untreated control culture, while most other glyphosate doses did not alter SCFA levels. We also measured *Enterobacteriaceae* levels as glyphosate is known to affect this taxonomic group by inhibiting the shikimate pathway. Overall, it was observed that glyphosate did not impact *Enterobacteriaceae* abundance at any of the concentrations tested (Supplementary Table 3).

Considering an average weight of a healthy infant of 3 years old around 15 kg, 20% of absorption of glyphosate at intestinal level, and assuming a steady-state concentration in the SHIME system, it was decided to select a concentration of 100 mg/L (14 mg glyphosate added to the system in each feeding cycle) for the long-term study, which would correspond to 1.2 mg/kg of body weight of glyphosate. This dose is in the range of the US ADI (1.75 mg/kg body weight/day).

### Microbial community activity during the long-term infant M-SHIME experiment

During the control period, SCFA levels were very stable within (on average 93.2% similar between consecutive time points in control period), and reproducible between both of the M-SHIME units (on average 91.7% similar), indicating stability and reproducibility of the experimental setup. This high stability guarantees that any effects observed during the treatment truly result from the administered test products, while the high reproducibility between each of the units allows direct comparison between the products on virtually identical microbial communities. No statistically significant differences were detected between the two different control weeks when microbial community activity was evaluated (Figure 1).

In contrast, large product-dependent differences were observed (Figure 2). The Roundup formulation always resulted in more alterations than glyphosate alone. While base consumption remained relatively unaffected in the ascending and transverse colon reactors upon glyphosate treatment (Figure 2A), base consumption was significantly enhanced by 55.0, 6.2 and 2.0 ml/day in the ascending, transverse and descending colon, respectively, upon addition of Roundup.

**Figure 2.**
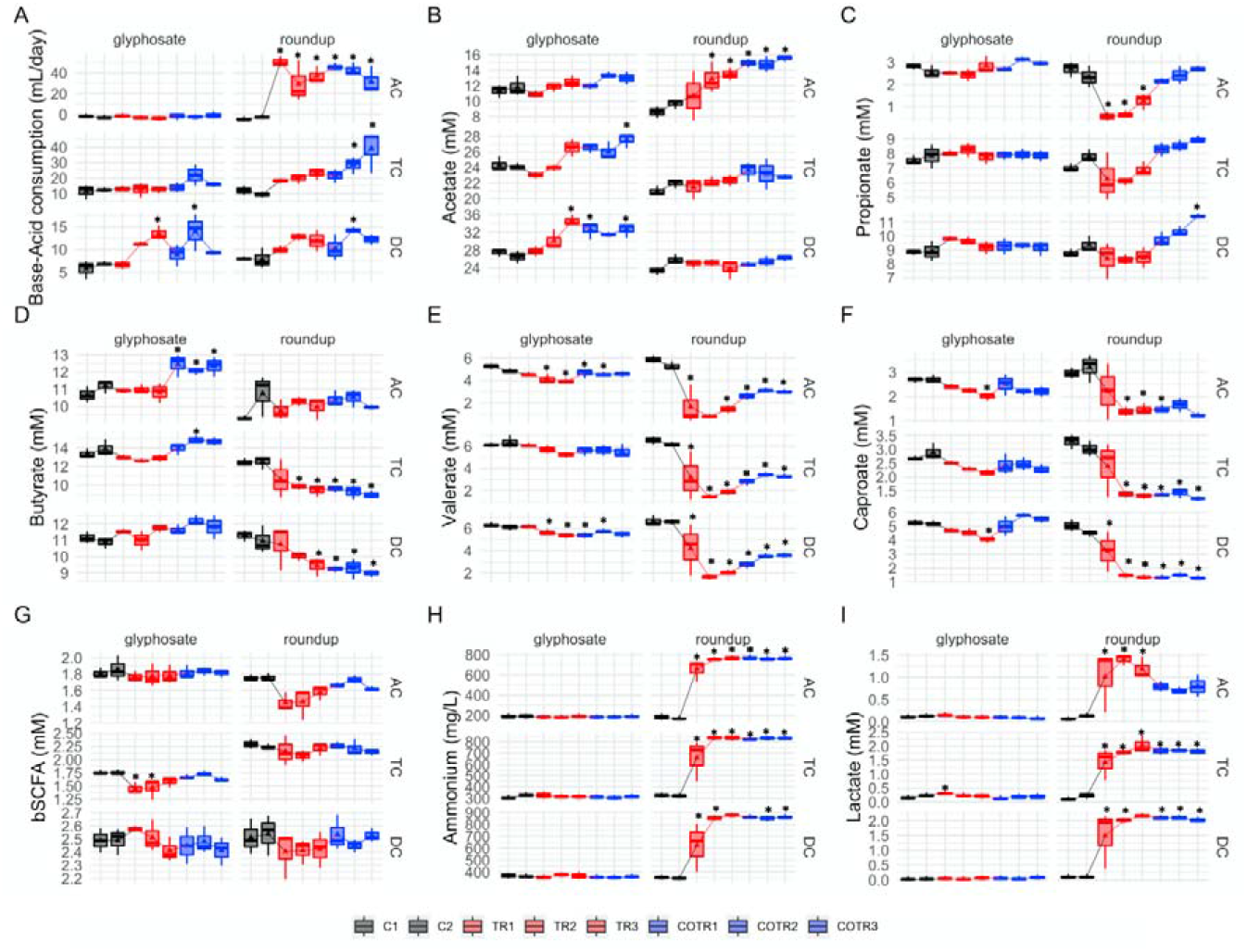
Analysis of microbial community activity after exposure to glyphosate or Roundup followed by co-exposure to the MegaSporeBiotic spore-based probiotic formulation in the microbiota derived from a healthy 3-year-old child. Average weekly base-acid consumption, SCFA levels (acetic acid, propionic acid, butyric acid, valeric and caproic acid), lactate, ammonium and branched SCFA (b-SCFA) in the culture chambers simulating the ascending (AC), transverse (TC) and descending (DC) colon during control (black), treatment (red) and co-treatment (blue) weeks. (*Indicates statistically significant differences relative to C1 after Tukey’s honestly significant difference (HSD) post hoc test with p<0.05; n = 3 per week).

Overall, glyphosate had limited effects on SCFA levels (Figure 2B to 2F) with acetate, propionate, butyrate, caproate and valerate levels remaining unaffected in the transverse colon, and only a slight increase in the amount of valerate (−1.09 mM and −0.74 mM for C2 vs TR3 in TC and DC, respectively) and caproate (−0.71 mM and −1.16 mM for C2 vs TR3 in TC and DC, respectively) was observed in the transverse and descendant colon areas. However, during the co-treatment period with MegaSporeBiotic prebiotic, an increase in butyrate of 1.63, 1.0 and −0.1 mM (from TR3 to COTR1) was observed in the ascending, transverse and descending colon, respectively (Figure 2D).

Differences in SCFA levels were larger after exposure to Roundup (Figure 2B to 2F). Roundup increased acetate production by 4.9 mM in the ascending colon, whereas acetate levels remained unaffected in the transverse and descendant colon during the treatment period. The SCFAs most affected by Roundup were propionate, valerate and caproate, which had their levels reduced in the ascending and transverse colon, with a similar trend being observed in the descending colon. For instance, Roundup caused a decrease of propionate levels by 1.8 mM, valerate levels by 3.9 mM and caproate levels by 1.75 mM in the ascending colon from the end of the control period (C2) to the end of the treatment period (TR3). Interestingly, addition of MegaSporeBiotic resulted in recovery of propionate levels in all colonic regions during Roundup treatment. In the distal colon areas (TC and DC), propionate levels even increased above levels observed during the control period (Figure 2C).

The human intestine harbours both lactate-producing and lactate-utilizing bacteria. Lactate produced by bacteria decreases the pH of the intestinal environment. Overall, treatment with glyphosate did not affect lactate levels in all colonic regions, except for a slight initial increase in the ascending and transverse colon reactors (Figure 2I). Roundup on the other hand significantly increased lactate levels in all colonic areas. The lactate levels were increased by 1 mM in the ascending colon, 1.8 mM in the transverse colon and 2 mM in the descending colon (comparison C2 to TR3). Roundup but not glyphosate significantly increased ammonium levels in all colon regions (Figure 2H). Ammonium levels were increased by 598 mg/L in the ascending colon, 518 mg/L in the transverse colon and 526 mg/L in the descending colon (comparison C2 to TR3).Overall, addition of MegaSporeBiotic did not affect ammonium levels during glyphosate and Roundup treatment.

### Global metabolomics

We next explored the changes in gut microbial activity in greater detail using a global metabolomics approach. A total of 842 biochemicals (721 compounds of known identity and 121 compounds of unknown structural identity) were detected. Concentrations of SCFA, which were measured quantitatively and detected in the global untargeted metabolomics, correlated well for caproate (R^2^ = 0.96, p-value < 2.2e-16), butyrate (R^2^ = 0.82, p-value = 8.0e-13) and valerate (R^2^ = 0.93, p-value < 2.2e-16), and confirmed the quality of our data (Supplementary Figure 1).

We used a multivariate strategy to understand the effects of glyphosate and Roundup in the different SHIME compartments. The metabolome changes were first visualized by plotting each sample in a space defined by the two principal components of a principal component analysis (PCA) (Figure 3A). The first component separated the group of samples by colon regions. The second component separated the samples exposed to the Roundup formulation from the other samples, whereas samples exposed to glyphosate did not separate from the control group. In order to understand which metabolites were driving these differences stemming from the treatment with Roundup, we built an OPLS-DA model on the basis of the PCA results. The OPLS-DA model separating the samples exposed to Roundup from the rest of the samples, appropriately classified all samples (R2X = 0.314, R2Y = 0.991, and Q2 = 0.93). The small difference between R2Y and Q2 (<0.2) and the Q2 value was greater than 50% revealed an excellent predictive capability. A 1000-time permutation test further validated the OPLS-DA model as the empirical p-values for R2Y (p = 0.001) and Q2 (p = 0.001) indicated that the observed effects of Roundup are not part of the distribution formed by permuted data. OPLS-DA models built by discriminating the samples according to their exposure to glyphosate, or to MegaSporeBiotic, were not found to be of sufficient quality to allow reliable conclusions and were thus not explored in greater detail.

**Figure 3.**
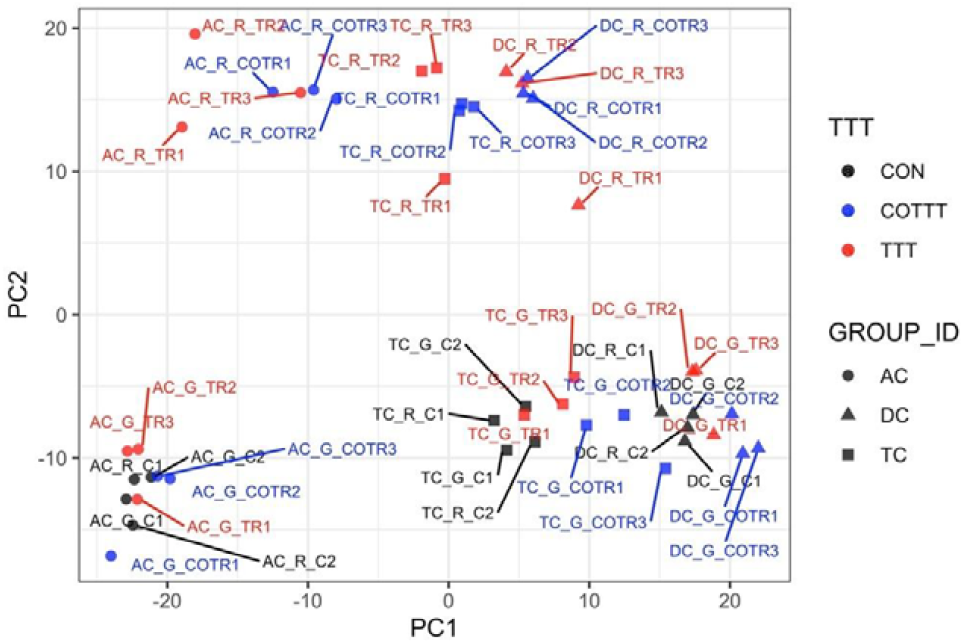
Principal component (PC) analysis to understand source of variation in the global metabolome profiles indicates that Roundup causes large scale disturbances in microbial activity. The metabolome changes are visualized by plotting each sample in a space defined by the two principal components of a PC analysis. The name of each sample is XX_Y_Z, and is providing indication on the metadata (XX corresponds to the ascending (AC), transverse (TC) and descending (DC) colon areas; Y to glyphosate (G) or Roundup treatment; Z to control (C1, C2, black), treatment (TR1, TR2, TR3, red) and co-treatment (COTR1, COTR2, COTR3, blue) weeks. TTT = treatment period of the infant M-TWIN SHIME.

We then evaluated which metabolites were affected by Roundup, and also performed a metabolic pathway enrichment analysis. The metabolites which are the best discriminators were selected using VIP scores. A selection of metabolites, which were found to be relevant for glyphosate toxicology is presented in Figures 3B to 3M, whilst plots for the 125 metabolites with a VIP score >1.5 are available as supplementary figure S8. Although glyphosate levels were unsurprisingly found to be the most variable (Figure 3C), the levels of this compound were affected only by the Roundup treatment. This could suggest that the surfactant co-formulants present in Roundup increased glyphosate availability. The most variable metabolite increased after treatment with Roundup was found to be one of unknown structural identity (Figure 3B), which suggests that glyphosate or Roundup metabolic pathways are still not fully elucidated. Levels of SCFAs (Figure 3F to 3I) were decreased confirming the results of the targeted analysis of microbial activity (Figure 1). Overall, most of the metabolites which were altered by Roundup exposure had their levels decreased, and were part of many microbial biochemical pathways, such as vitamin or hormone precursors (Figure 3J to 3L), which suggests that this pesticide mixture exerts global inhibitory effects on microbial metabolism. Methylphosphate (Figure 3D) was among the few metabolites having its levels increased by Roundup exposure. In addition, we noticed the solanidine had its levels decreased by both Roundup and glyphosate exposure (Figure 3A). Statistical significance of pathway enrichment was tested using a two-sided hypergeometric test. This revealed that Roundup affected the pathways for long chain polyunsaturated fatty acid (n3 and n6) metabolism (p = 0.00003), short chain fatty acid metabolism (p = 0.02), (hypo)xanthine/inosine containing purine metabolism (p = 0.04), and xenobiotic chemical metabolism (p = 0.003). While most of the SCFA fatty acids had their levels decreased, polyunsaturated fatty acid (n3 and n6) were among the small proportion of metabolites which had their abundance increased by Roundup exposure, including docosapentaenoate (22:5n3 and 22:5n6), docosahexaenoate (22:6n3), eicosapentaenoate (20:5n3), dihomo-linolenate (20:3n3 and 20:2n6), linoleate (18:2n6), arachidonate (20:4n6) and cis-4-decenoate (10:1n6) (Supplementary Figures S8).

### Microbial community composition during the long-term M-SHIME culture

Previous studies have reported that glyphosate and its formulations exerted a wide spectrum effects on bacterial growth and affected gut microbial community composition at the phylum level, in particular for Bacteroidetes phylum and Firmicutes phylum. In order to evaluate whether we can identify which bacterial species are affected by glyphosate and Roundup, we sequenced amplicons of the V3-V4 regions from the 16S rRNA genes.

In the present study, qPCR was also used to monitor the abundance of taxa with known health roles. This analysis was done both in the luminal and the mucosal gut microbiota populations. The results from the qPCR experiments for the Bacteroidetes phylum and Firmicutes phyla (Supplementary Figure 3) as well as for *Akkermansia muciniphila, Faecalibacterium prausnitzii, Bifidobacterium spp., Lactobacillus spp*. (Supplementary Figure 4), presented a substantial degree of technical variation, which limited the conclusiveness of this compositional analysis. This is reflected by the variations exhibited during the control period, with large changes in bacterial abundance observed between C1 and C2. In addition, poor correlations between the results of the qPCR and the 16S RNA gene analysis for major taxa such as Firmicutes and Bacteroidetes can be observed (Supplementary Figure 2). Thus, our analysis of bacterial composition only provided preliminary evidence, which would need to be confirmed in other studies. It is however noteworthy that there is a trend towards an increase in *Lactobacillus spp*. levels caused by the exposure to glyphosate and Roundup in both the luminal and the mucosal compartment (Supplementary Figure 5A). Lactobacilli are regarded as beneficial saccharolytic bacteria that are capable of producing high concentrations of lactate. The increase in lactobacilli could thus explain the increase in lactate levels in all colon compartments upon Roundup and glyphosate treatments (Figure 3 and 4).

**Figure 4.**
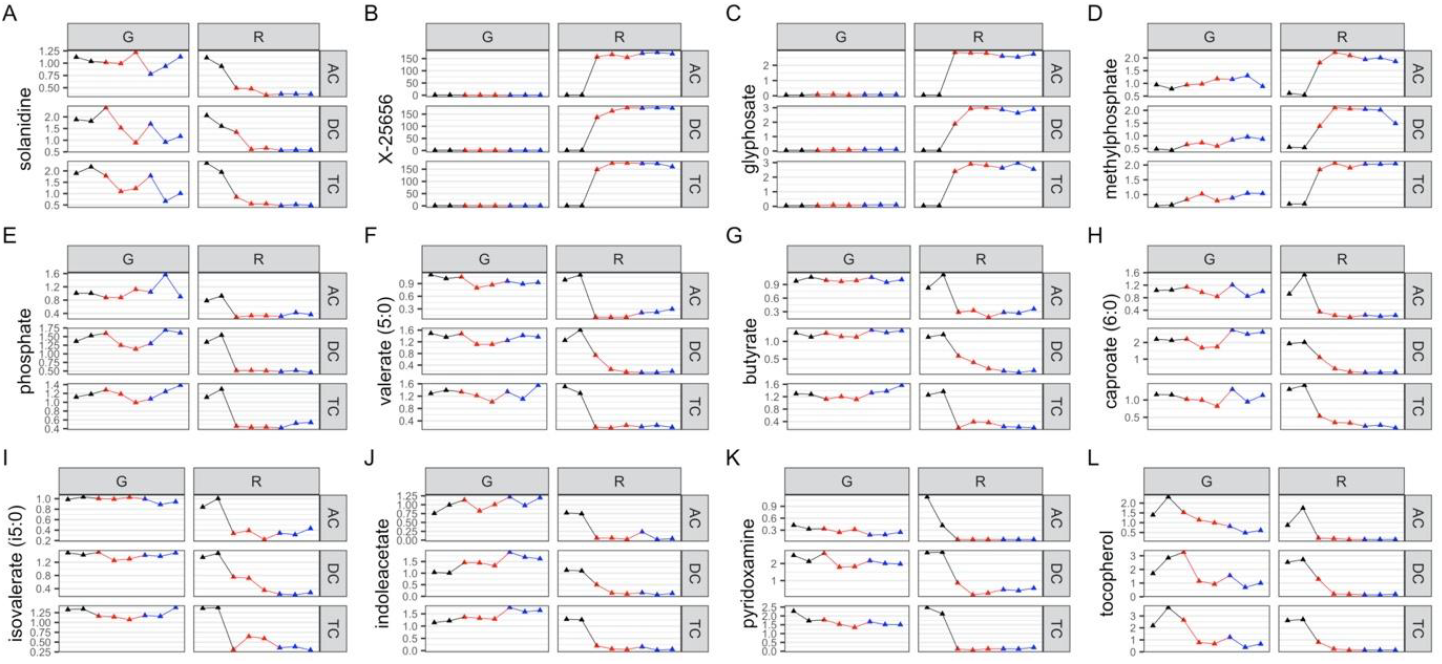
Variation in abundance of metabolites mostly affected by Roundup exposure. Normalised abundance levels are presented for key metabolites in glyphosate (G) and Roundup (R) SHIME in the ascending (AC), transverse (TC) and descending colon (DC) during control (black), treatment (red) and co-treatment (blue) weeks, among 125 metabolites discriminating the Roundup exposed samples in an OPLS-DA models (Supplementary Figures S8).

The Firmicutes-to-Bacteroidetes ratio (F/B) has been proposed as a potential biomarker of gut dysbiosis associated with specific diseased conditions (Magne et al. 2020). Glyphosate significantly reduced F/B in the ascending and distal colon in the luminal compartment (p<0.05), a trend also observed for the mucus environment in all the compartments (**Figure XXX**). Contrastingly, Roundup treatment increased F/B in the luminal (DC) and mucosal (AC and TC) compartments (p<0.05) (Supplementary table S4). Cotreatment with the spore-based probiotic MegaSporeBiotic did not rescue F/B ratios.

Microbiota DNA sequencing provided an average of 112,230 reads for each sample (min 57,313 - max 139,760). A summary of the composition for the different compartments during the control period indicated that the gut microbiome was dominated by Firmicutes and Bacteroidetes. The phylum Verrucomicrobia accounting for approximately 5% of the assigned abundance was mainly represented by *Akkermansia muciniphila*. This taxonomic profile is typical of people from Western societies. At lower taxonomic levels, we observed that a total of 3 genera accounted for approximately ~70% of the assigned abundance (Figure 5A). This included the *Megasphaera* spp., which are generally not among the most abundant species in the general population from Western societies, but which is abundant in the gut microbiome from individuals whose diet is rich in dairy products, which can be the case for young children (Dhakan et al. 2019).

**Figure 5.**
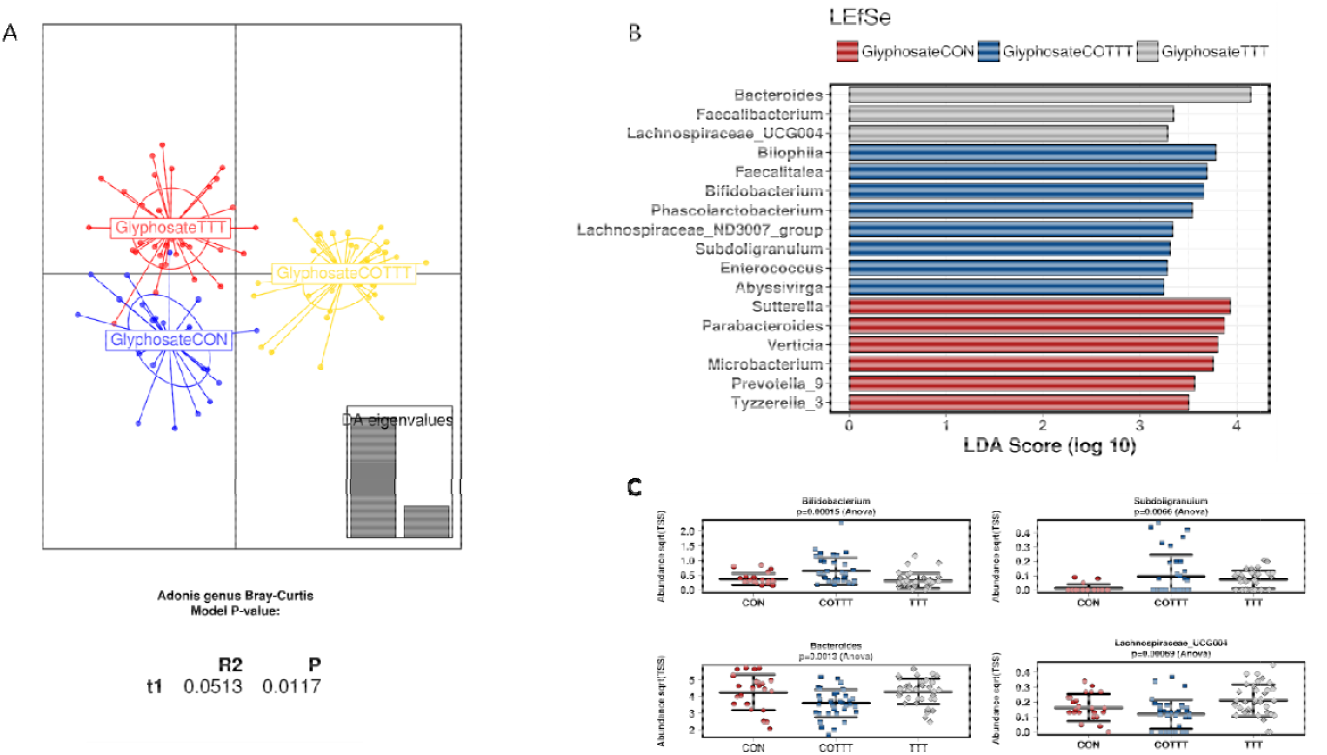
Effect of glyphosate on infant gut microbial structure. (**A**) Discriminant analysis of principal components and Adonis test based on Bray-curtis distance at genus level. (**B**) Linear discriminant analysis Effect Size for the glyphosate arm during control (CON), treatment (TTT) and co-treatment (COTT) conditions. (**C**) Strip char plots of selected features significantly different (ANOVA, p<0.05) between conditions.

Analysing glyphosate and Roundup treatments together, the DAPC plot showed a different clustering based on control, treatment or co-treatment (p <0.05, Adonis test at genus level) (Supplementary Figure S6A). The only family affected by treatments was *Lactobacillaceae*, due to an enrichment in glyphosate or Roundup treated reactors (LDA > 3), whereas co-treatment enriched *Synergistaceae, Desulfovibrionaceae, Acidaminococcaceae* and *Bifidobacteriaceae* (LDA > 3) among others (Supplementary Figure S6B). To define the effect of specific treatments on infant gut microbiota, samples from glyphosate or Roundup conditions were independently analysed. Samples from glyphosate treatment and further cotreatment with MegaSporeBiotic probiotic clustered separately from the control (p < 0.05, Adonis at genus level) (Figure 5A). Samples from the control condition were enriched in *Sutterella, Parabacteroides, Verticia, Microbacterium, Prevotella* and *Tyzerella* genera (LDA > 3). During the glyphosate treatment, an enrichment in *Bacteroides, Faecalibacterium* and *Lachnospiraceae_*UCG004 was observed. Finally, probiotic treatment enriched *Bilophila, Faecalitalea, Bifidobacterium, Subdoligranulum* and *Phascolarctobacterium*, among other genera (Figure 5B, 5C).

Roundup treatment also induced a different clustering from control and co-treatment (p < 0.0001, Adonis at genus level) (Figure 6A), with enrichment in *Bacteroides, Faecalitalea, Roseburia, Hydrogenoanaerobacterium*, and *Faecalibacterium*. MegaSporeBiotic probiotic co-treatment enriched, in addition to some genera described for the glyphosate condition, *Akkermansia* and *Cloacibacillus*. When analysing the data using a non-metric multidimensional scaling (NMDS) of Bray-Curtis distances (Supplementary Figure S7), no clear clustering based on treatment was observed, suggesting a potential effect of time on composition profiles.

**Figure 6.**
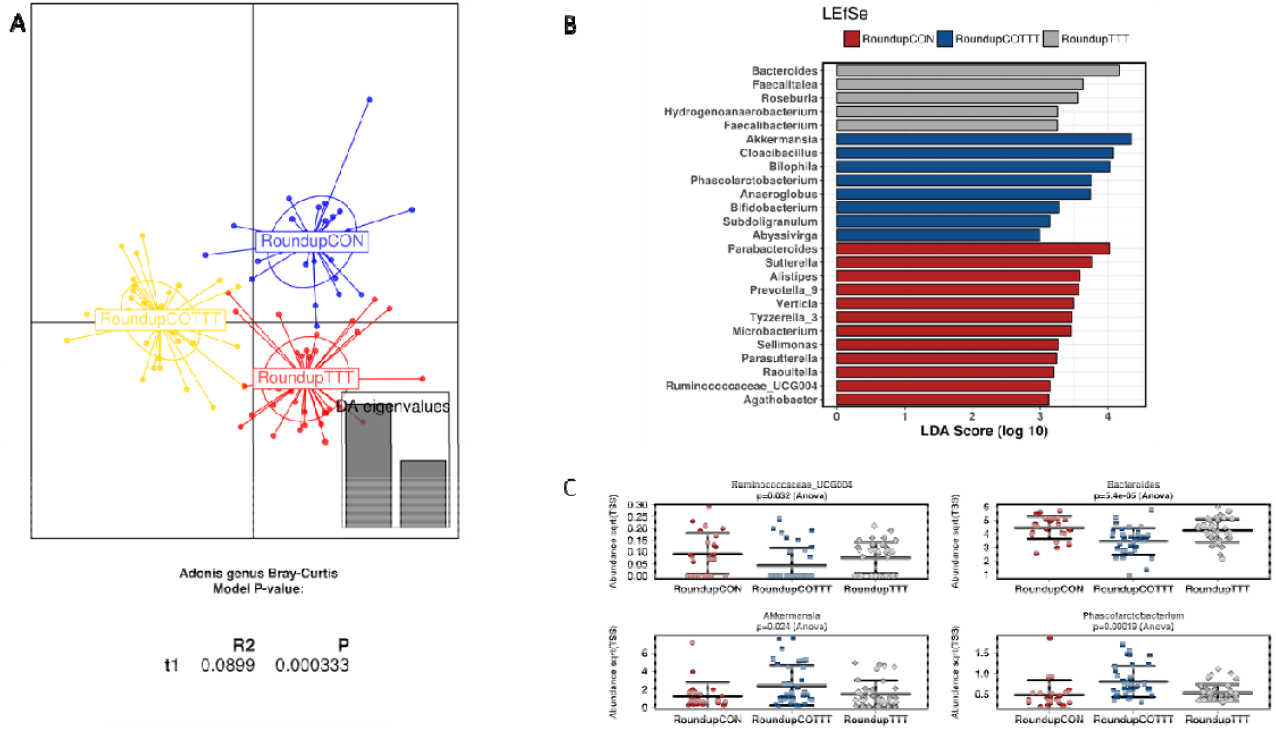
Effect of Roundup on infant gut microbial structure. (**A**) Discriminant analysis of principal components and Adonis test based on Bray-curtis distance at genus level. (**B**) Linear discriminant analysis Effect Size for the Roundup arm during control (CON), treatment (TTT) and co-treatment (COTT) conditions. (**C**) Strip char plots of selected features significantly different (ANOVA, p<0.05) between conditions.

Glyphosate and Roundup treatments were further compared, showing increased levels of *Pseudomonadaceae, Collinsella, Phascolarctobacterium, Sutterella* and *Lachnospiraceae*__ND3007_group in the presence of Roundup (LEfSe, LDA > 2, Figure 7A; heat tree based on non-parametric Wilcoxon Rank Sum test, Figure 7B). At a lower phylogenetic level, OTU37 (*Bacteroides* spp.), OTU82 (*Klebsiella* spp.) and OTU88 (*Collinsella* spp.) were significantly different between glyphosate and Roundup treated cultures (Figure 7C, zero-inflated Gaussian fit mixed model MicrobiomeAnalyst software).

**Figure 7.**
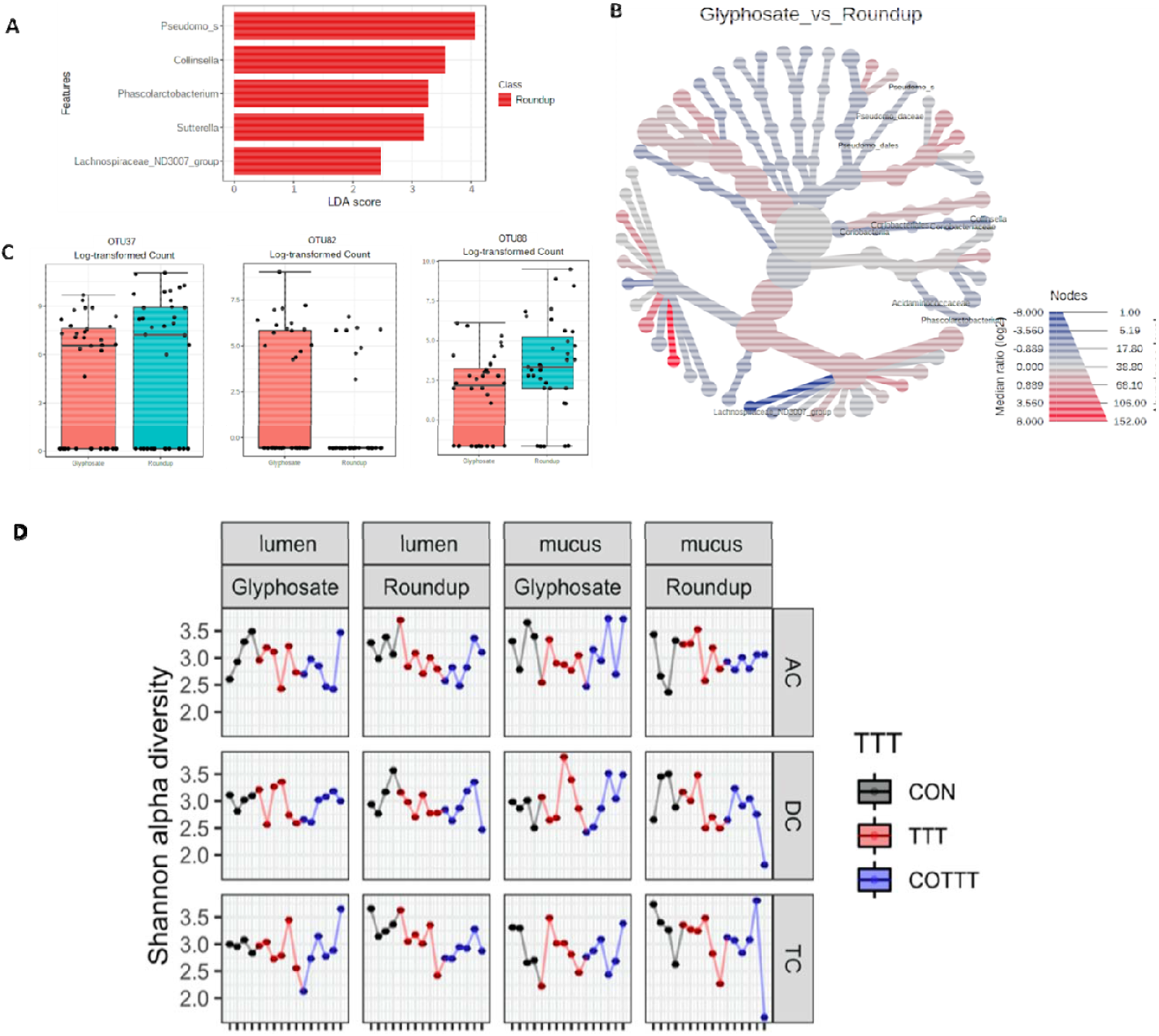
A 16S rRNA gene amplicon sequencing approach to assess alterations in gut microbiome composition. (**A**) Linear discriminant analysis Effect Size including treatment samples from glyphosate and Roundup treatment arms of the study. (**B**) Heat tree analysis of differences between glyphosate and Roundup treatment at genus level. Significantly different taxa between groups are written in the figure.

We found a statistically significant difference in average bacterial Shannon alpha diversity (Figure 7C) by the treatments (p = 0.02), but it was not different between glyphosate and Roundup (p = 0.42), between the mucus and the lumen (p = 0.87), or between the different SHIME compartments (p = 0.64). Tukey multiple comparisons of means further showed that alpha diversity decreased with the pesticide treatment (p = 0.03), and later recovered after exposure to MegaSporeBiotic (Figure 7C).

As for the analysis employing qPCR, the changes in bacterial composition derived from 16S rRNA gene sequencing presented a substantial degree of technical variation reflected by the variations exhibited during the control period as illustrated for the most frequently found genera *Megasphaera*, *Bacteroides*, *Lachnoclostridium* and *Akkermansia* (Supplementary Figure 5).

Altogether, our study of the changes in gut microbial activity and composition after a long-term exposure to glyphosate and Roundup reveals profound changes in the fermentation activity and metabolite profiles, while changes in composition profiles were less pronounced with the current experimental design.

## DISCUSSION

We describe for the first time the effects of glyphosate and a formulated glyphosate Roundup herbicide on human gut microbiota. Using an in vitro technology mimicking the entire gastrointestinal tract revealed that glyphosate caused large scale disturbances in the activity of gut microbiota obtained from a healthy infant.

Both glyphosate and Roundup caused a change in the fermentation activity of the infant gut microbiota as reflected by the increase in base consumption. This is because optimal environmental conditions are maintained in the SHIME by base addition in order to maintain the pH, which can change as a result of bacterial fermentation. This acidification of the gut microbial ecosystem by glyphosate can be linked to an increase in the production of acetate and lactate, as bacterial production of these compounds tends to acidify the medium. This in turn favours the growth of bacteria, which can tolerate a lower pH such as lactobacilli which had their growth increased during the treatment period in this study. Despite the fact that specific *Lactobacillus* and *Bifidobacterium* strains are currently accepted as probiotics (Hill et al. 2014), composition of mucosal-associated microbes in inflammatory bowel patients has shown increased proportions of Bifidobacteria and lactobacilli, against a decrease in butyrate-producing bacteria (Wang et al. 2014). This study suggests that imbalance between gut commensals, might influence host inflammatory responses. This also correlates well with findings of our recent laboratory animal study showing that rats exposed prenatally until adulthood to glyphosate presented increased levels of lactobacilli (Mesnage et al. 2021a).

The pH of the gut environment is key in maintaining ecosystem homeostasis and it has been described as the strongest driver for microbial community structure *in vitro* (Ilhan Zehra et al.). When the pH of the gut microbiome environment decreases, it impairs the growth of pH sensitive bacteria such as *Bacteroides* spp (Walker et al. 2005) and inhibits lactate-consuming species, leading to lactate accumulation (Ilhan Zehra et al.). Lactate accumulation has been linked to butyrate and propionate reduction, due to major shifts in microbiota composition, with Bacteroidetes and anaerobic Firmicutes being substituted by Actinobacteria, lactobacilli and Proteobacteria (Ilhan Zehra et al.). Since some members of the Bacteroides family are known propionate or butyrate producers (Walker et al. 2005), the acidification of the gut microbial environment by glyphosate is also likely to explain the decrease in specific bacterial metabolites. This is also visible in our recent animal study in which Bacteroidota abundance was decreased by glyphosate exposure (Mesnage et al. 2021a). In addition, reductions in butyrate and propionate induced by pesticides can be potentially caused by alterations in enzymatic paths involved in cross-feeding interactions, for example involving co-A transferase (Louis and Flint 2017) or the acrylate pathway.

Microbial SCFAs are essential for maintaining gut homeostasis through different mechanisms involving control of gut barrier integrity, regulating the luminal pH, mucus production, providing energy for epithelial cells and modulating mucosal immune function (Blaak et al. 2020). Lactate accumulation has been described in colitis patients (Vernia et al. 1988) and it can also be used as a growth substrate by sulfate-reducing gut bacteria (Marquet et al. 2009), promoting the formation of toxic concentrations of hydrogen sulphide. Reduction of butyrate and propionate production and lactate accumulation induced by a glyphosate-based herbicide indicates functional dysbiosis in our *in vitro* SHIME system. However, potential consequences on the host still require further investigation.

Both the production of ammonium and branched-SCFA result from protein degradation and reflect proteolytic activity of the gut microbiota. Ammonium production was substantially increased by Roundup exposure but not by glyphosate. This could be associated with the increased abundance of lactobacilli as the higher metabolic activity of this class of bacteria can enhance deamination processes, which is the main pathway of amino acid fermentation in humans. It is worth noting that an increased production of ammonium is detrimental because it can activate cell proliferation mechanisms in colonocytes and subsequently promote colon carcinogenesis (Visek 1978).

Global metabolomics revealed disturbances in a large number of metabolic pathways. However, the relevance to health of most of these changes are still elusive. This is the case for solanidine levels, which were decreased by both Roundup and glyphosate exposure. This is particularly interesting because this has already been described in our study involving subchronic exposure to another Roundup formulation in the rat gut microbiome (Mesnage et al. 2021b), and in potato plants grown in glyphosate-treated soil (Rainio et al. 2020). Methylphosphate was among the few metabolites, which had its abundance stimulated by Roundup. Since glyphosate is an organophosphate compound, this observation may originate from previously unknown biotransformation of glyphosate. Perhaps the most surprising metabolome changes we observed was the increase in levels of n3 and n6 long chain polyunsaturated fatty acids (PUFA) Some limited groups of bacteria can synthetize distinct unsaturated fatty acids such as arachidonic acid (20:4 n-6), eicosapentaenoic acid (20:5 n-3), and docosahexaenoic acid (22:6 n-3) (Okuyama et al. 2007; Yoshida et al. 2016). Anaerobic *de novo* synthesis of PUFA is mediated by the PUFA synthase complex, but only described in environmental microorganisms up to now (Metz James et al. 2001; Okuyama et al. 2007; Yoshida et al. 2016). On the other hand, human gut microbiota modulate PUFA metabolism, involving multiple microbial species (Ewaschuk et al. 2006; McIntosh et al. 2009). Recently, docosahexaenoic acid, and arachidonic acid were positively correlated with *Bacteroides* in patients with chronic spontaneous urticaria (Wang et al. 2020). It is also possible that PUFA are formed from the metabolism of Roundup surfactants, which are in general synthetized from vegetable oils or animal fat (Mesnage et al. 2019). However, surfactants employed in formulating Roundup herbicides are generally considered as confidential business information by the manufacturer and thus not generally disclosed (Mesnage et al. 2019). Therefore, it is not fully clear if the blend of surfactant entering in the composition of Roundup PROMAX used in this investigation could contain surfactants synthetized from PUFA-containing oils.

The increase of long-chain fatty acids has been positively correlated with an elevated abundance of *Lactobacillus* in a rodent model of neonatal maternal separation characterized by accelerated colonic motility and gut dysbiosis, suggesting a relevant role of these molecules on host-microbiota interplay and potentially affecting host metabolism (Zhao et al. 2018). Fatty acid profiles in aquatic ecosystems have been proposed as biomarkers of pesticide exposure (Gonçalves et al. 2021). Short-term exposure to glyphosate or Roundup in the sea cucumber *Holothuria forskali* perturbed fatty acid composition, including some essential fatty acids (Telahigue et al. 2021). Changes in composition of PUFAs; i.e., alteration in the n3/n6 ratio, have been associated with increased incidence and prevalence of being overweight and obesity, linked to inflammatory responses (Jovanovic et al. 2021). Altogether, we propose pesticide disruption of fatty acid human gut metabolome as a potential biomarker of exposure, and also a mechanism of metabolic disturbance potentially influencing the host.

Whereas structural changes based on 16S rRNA gene sequencing did not indicate dramatic shifts on microbial composition, alpha diversity was reduced by exposure to both glyphosate and Roundup. Lower alpha diversity indices during infancy have been correlated with lower cognitive performance (Carlson et al. 2018), and negative health outcomes, including type 1 diabetes (Kostic et al. 2015) and asthma (Abrahamsson et al. 2014). The dynamic alterations in functional capacity of the simulated infant gut ecosystem suggest an impact of glyphosate or Roundup exposure on host function. A study in rats assessing prenatal exposure to glyphosate or two Roundup formulations showed the gut microbiome of F1 offspring was affected by both treatments, with a reduction in Bacteroidota abundance, concomitant to increased levels of Firmicutes and Actinobacteria (Mesnage et al. 2021a).

Within the temporal framework used in this study, we observed a time-course modulation of the microbial metabolic landscape suggesting structural resilience of the infant gut ecosystem to glyphosate or Roundup exposure, but a functional dysbiosis affecting key microbial metabolites for microbiota-host cross-talk. For example, carboxyethyl-gamma aminobutyric acid (GABA), was significantly reduced in the ascending colon after Roundup treatment. GABA is linked to glutamate metabolism and the gut-brain axis and different microbial taxa regulate the GABAergic system in the human gut, including *Bifidobacterium* spp. (Duranti et al. 2020) and *Bacteroides* spp (Otaru et al. 2021).

Our results suggest that the addition of MegaSporeBiotic spore-based probiotic affected the gut microbiota and mitigated glyphosate-induced changes. Some Bacillus species are adapted to survive in the intestine (Cartman et al. 2008; Tam et al. 2006). In such cases, spores ingested with food are able to survive transit through the stomach after which they germinate and proliferate, a phenomenon that has been proven using *in vivo* studies (Cartman et al. 2008; Tam et al. 2006). Following growth and proliferation they re-sporulate as the bacteria pass through the intestine and are shed into the environment. Such germination and re-sporulation would be a necessity for certain health benefits, as was shown for the stimulation of the development of the gut-associated lymphoid tissue in rabbits by *B. subtilis* (Rhee et al. 2004). However, our study does not distinguish as to whether the Bacillus spores mitigated the effects of glyphosate by directly counteracting its effects, or if they add effects which are independent of glyphosate presence. Caution is also needed when extrapolating these findings to real-world glyphosate exposure. Our results and previous studies in animal model systems suggest structural and metabolic shifts in the gut environment due to exposure to pesticide residues (Tsiaoussis et al. 2019). The effect of these changes on human health remain to be determined. In addition, the conclusiveness of this study is very limited by its experimental design since it is very difficult to generalise conclusions from a study performed with one just biological sample. In addition, it is likely that the statistical power did not allow the detection of effects for taxa present a low abundance (less than 5%) in the gut microbiome samples. Further experiments will be needed to validate these first observations.

## Acknowledgements

This work was funded by Microbiome Labs (USA) and by the Sustainable Food Alliance (USA) whose support are gratefully acknowledged.

## SUPPLEMENTARY FIGURES AND TABLES

**Supplementary Figure 1.**
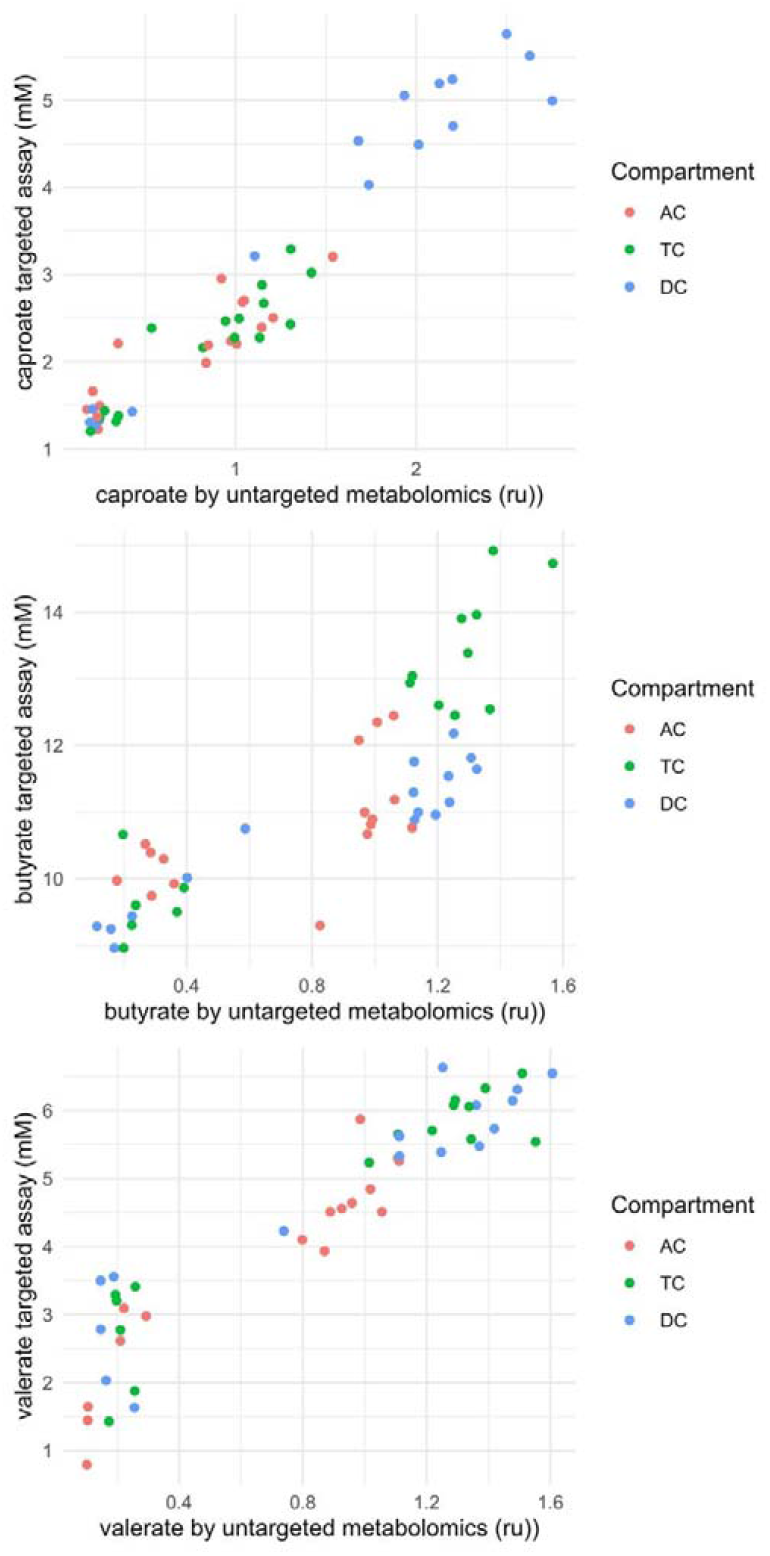
Absolute SCFA concentrations correlate to their relative abundance measured by untargeted metabolomics. Concentrations of SCFA caproate, butyrate and valerate correlate well with their relative abundance by global untargeted metabolomics in the ascending (AC), transverse (TC) and descending (DC) colon areas of the SHIME system.

**Supplementary Figure 2.**
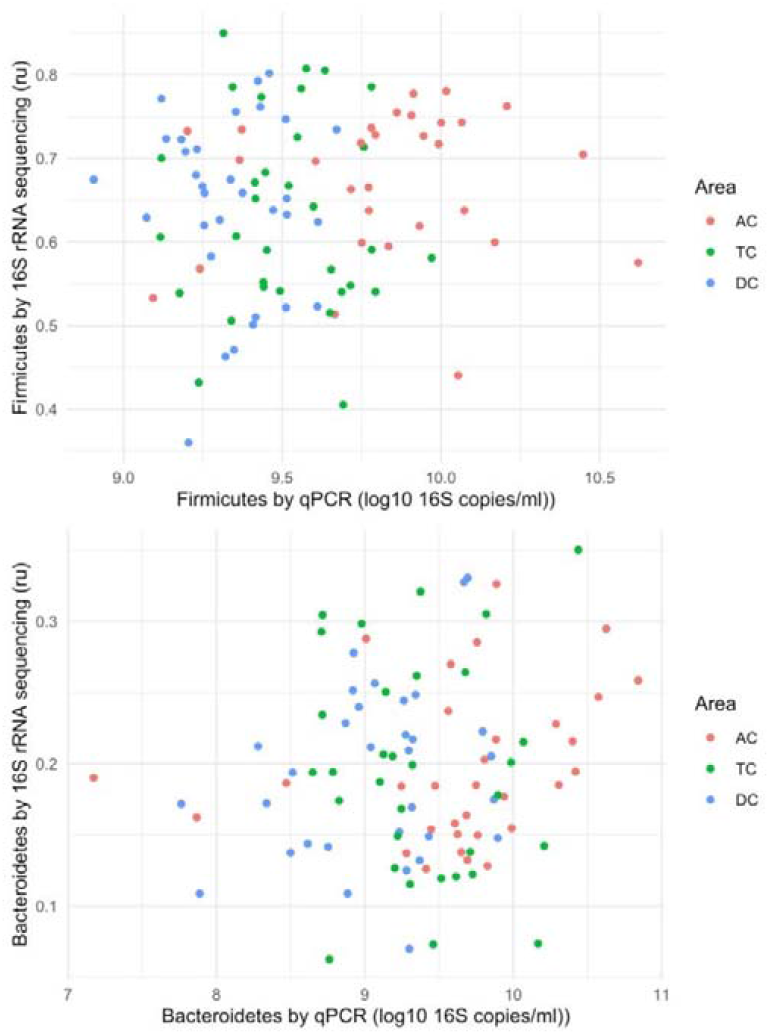
Bacteroidetes and Firmicutes levels poorly correlate between the qPCR and the 16S RNA gene sequencing results.

**Supplementary Figure 3.**
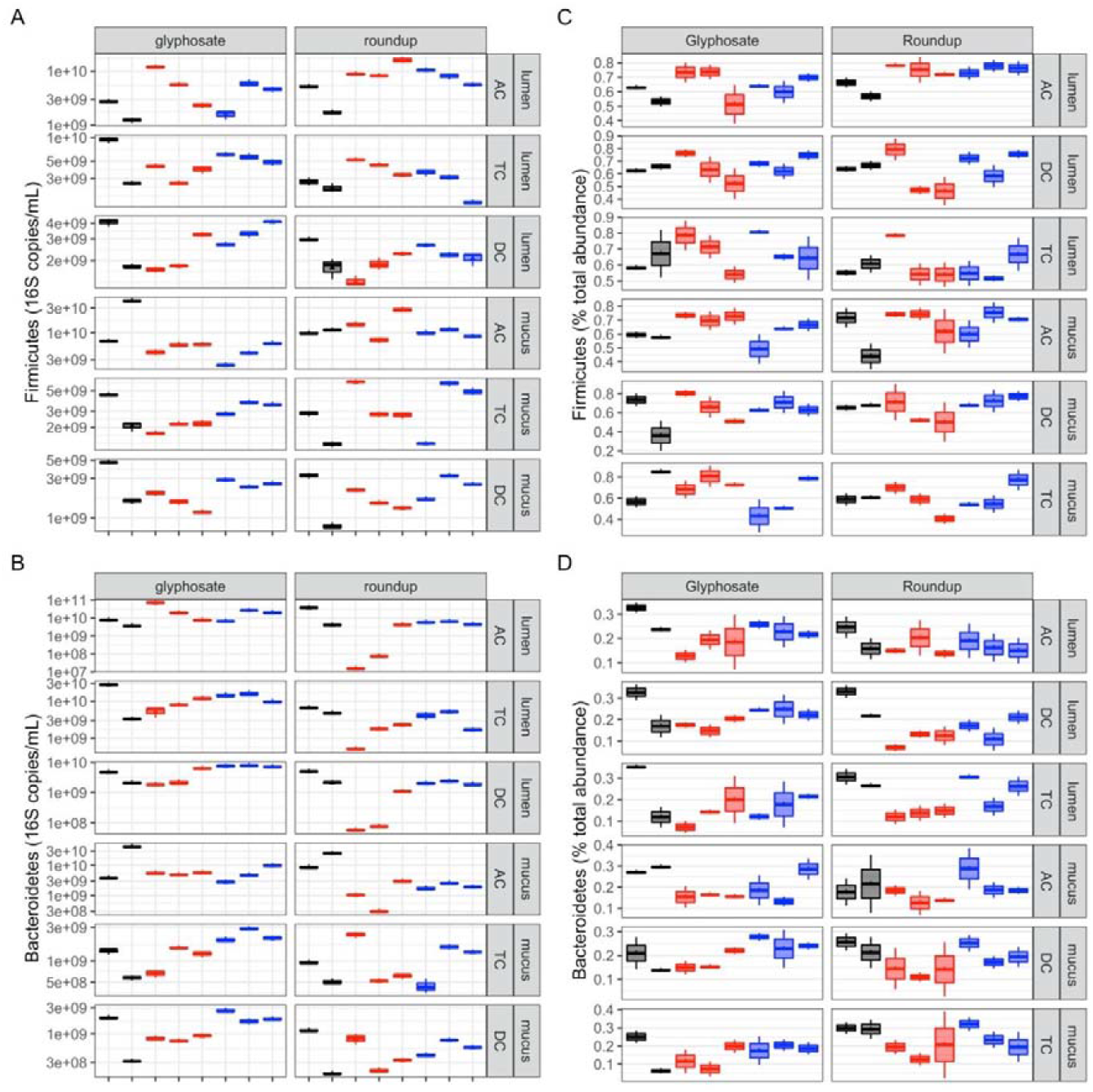
Analysis of the effect of glyphosate and Roundup on Firmicutes and Bacteroidetes levels. A qPCR analysis was used to measure the number of 16S RNA gene copies per ml for the two major phyla Firmicutes (**A**) and Bacteroidetes (**C**), and the results were compared to those obtained by 16S rRNA gene sequencing for the same taxa of Firmicutes (**B**) and Bacteroidetes (**D**) in the ascending (AC), transverse (TC) and descending (DC) colon compartments. Average weekly levels during control (C1-C2, black), treatment (TR1-TR3, red) and co-treatment (COTR1-COTR3, blue) weeks are shown.

**Supplementary Figure 4.**
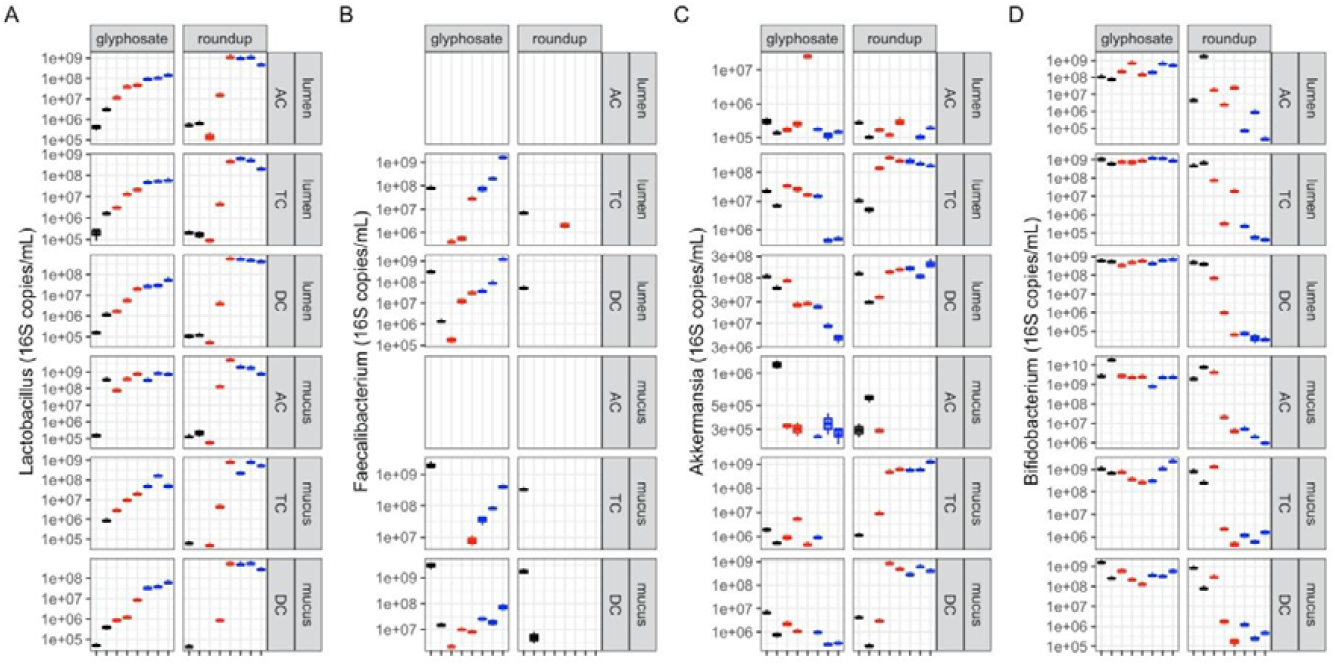
Analysis of the effects of glyphosate and Roundup on ***Lactobacillus spp, Faecalibacterium prausnitzii, Akkermansia muciniphila, Bifidobacterium spp***., levels by qPCR. A qPCR analysis was used to measure the number of 16S gene copies per ml for bacterial species with known roles in health, *Lactobacillus spp, Faecalibacterium prausnitzii. Akkermansia muciniphila, Bifidobacterium spp*., in the ascending (AC), transverse (TC) and descending (DC) colon compartments. Average weekly levels during control (C1-C2, black), treatment (TR1-TR3, red) and co-treatment (COTR1-COTR3, blue) weeks are reported.

**Supplementary Figure 5.**
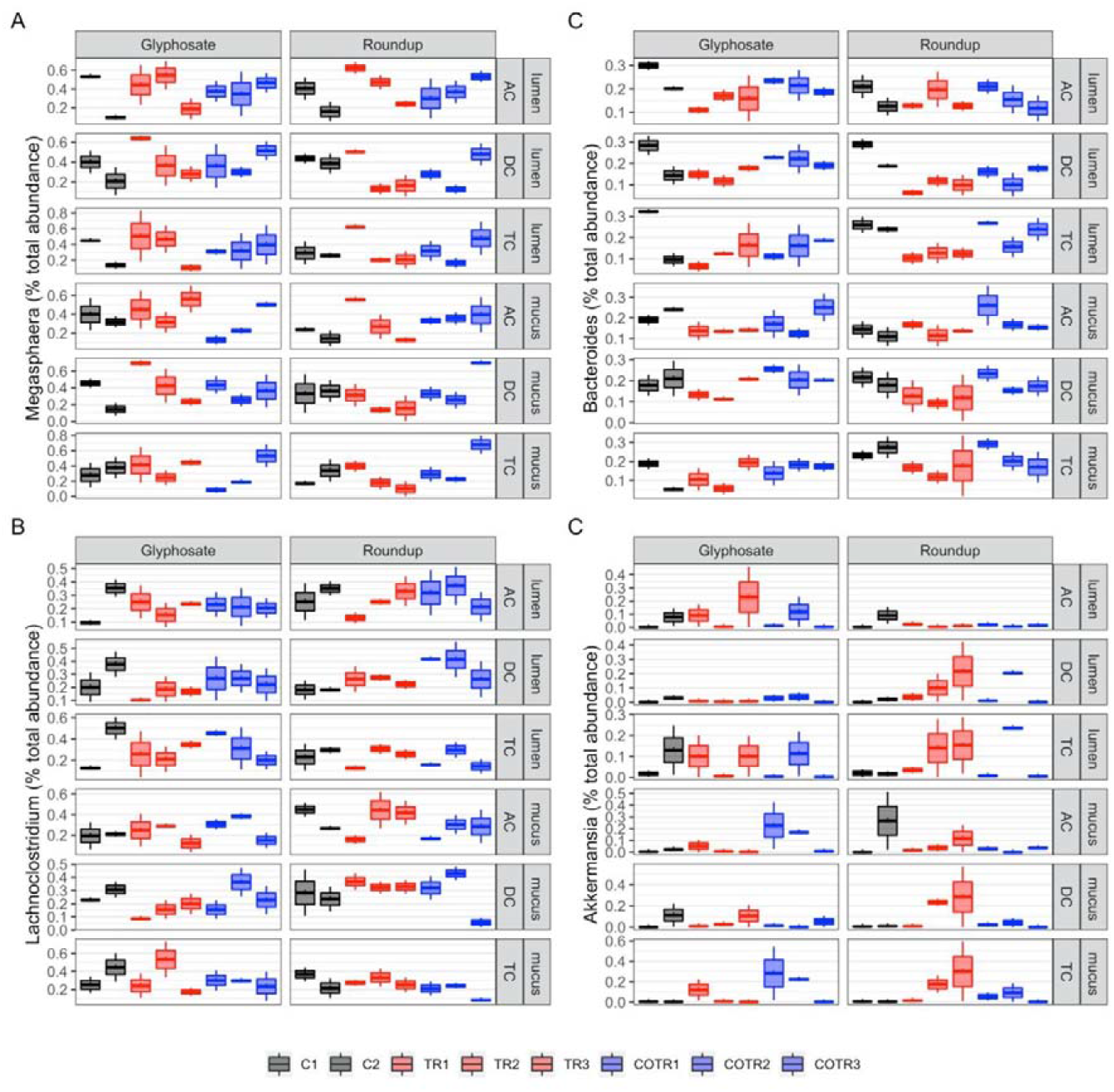
Analysis of the effects of glyphosate and Roundup on the 4 most frequent bacterial genera identified by 16S rRNA gene sequencing. Effects of the treatments are represented for Megasphaera (**A**), Bacteroides (**B**), Lachnoclostridium (**C**) and Akkermansia (**D**) in the ascending (AC), transverse (TC) and descending (DC) colon simulation chambers. Average weekly levels during control (C1-C2, black), treatment (TR1-TR3, red) and co-treatment (COTR1-COTR3, blue) weeks are reported.

**Supplementary Figure S6.**
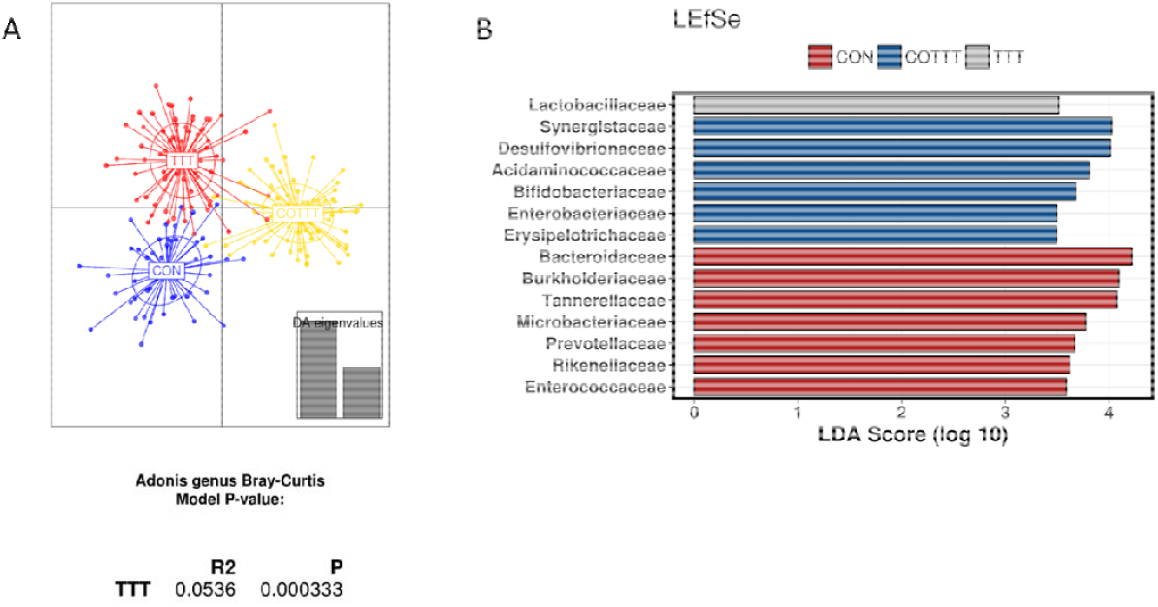
Effect of control, treatment and co-treatment conditions on microbial strucutre. (**A**) Discriminant analysis of principal components and Adonis test based on Bray-Curtis distance at genus level. (**B**) Linear discriminant analysis Effect Size considering control, treatment (glyphosate and Roundup), and co-treatment (glyphosate and Roundup + MegaSporeBiotic probiotic) period.

**Supplementary Figure S7.**
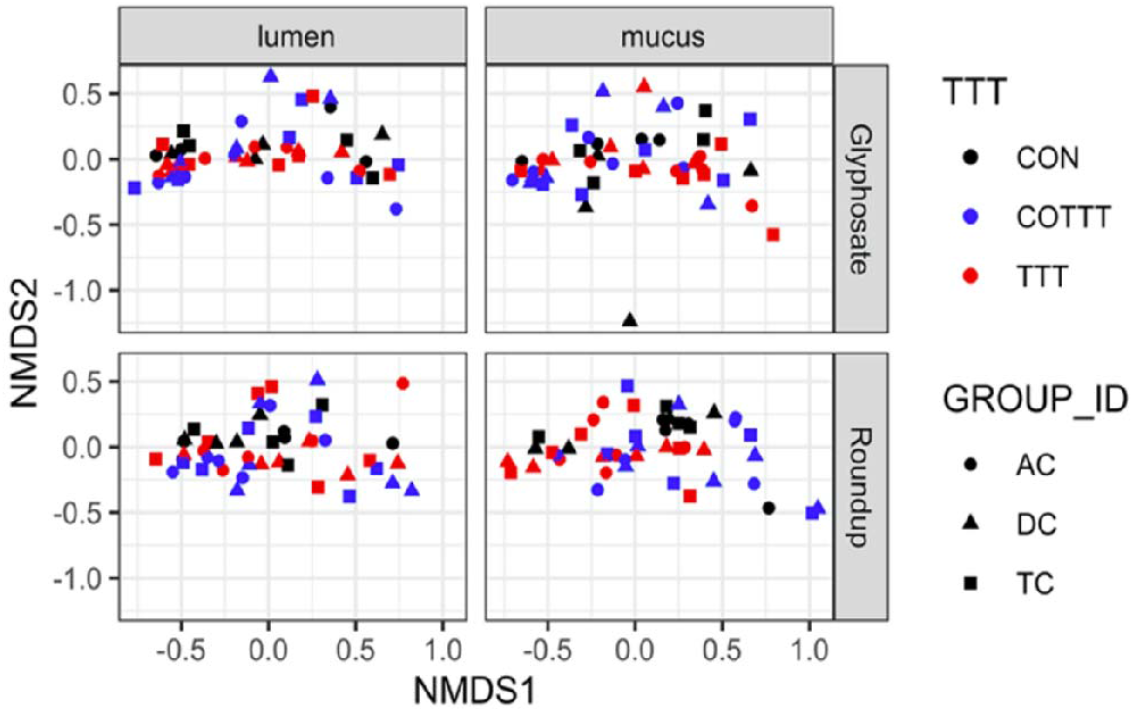
**Non-metric multi-dimensional scaling including treatment conditions, luminal and mucus compartment and colonic regions. AC = ascending colon, TC = transverse colon, DC = descending colon.**

**Supplementary Table S1.**
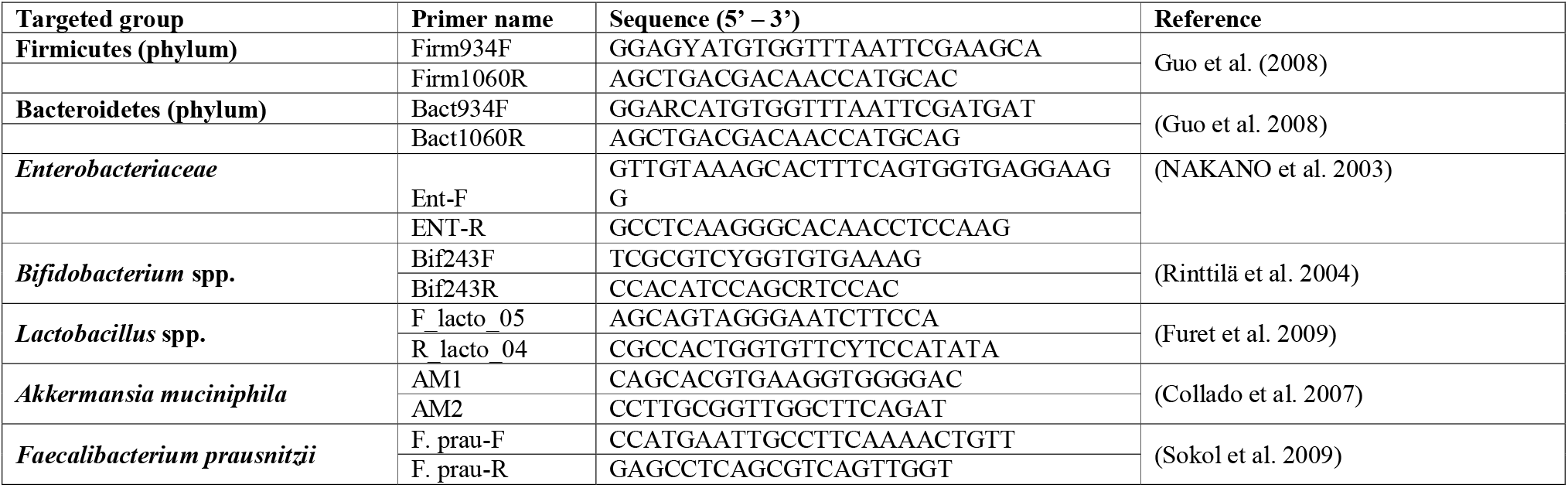
Primers used to quantify Firmicutes, Bacteroidetes, Bifidobacterium spp., Lactobacillus spp., Enterobacteriaceae, Akkermansia muciniphila and Faecalibacterium prausnitzii by qPCR.

**Supplementary Table S2.** qPCR conditions.

| Program | Cycle | Temperature (°C) | Heating (hh:mm:ss) | Ramp (°C/s) |
| --- | --- | --- | --- | --- |
| Pre-incubation | 1 | 95 | 00:10:00 | 1.6 |
| Amplification | 40 | 95 | 00:00:15 | 1.6 |
|  |  | 60 | 00:00:30 | 1.55 |
|  |  | 72 | 00:00:30 | 1.6 |
| Melting curves | 1 | 95 | 00:00:15 | 1.6 |
|  |  | 60 | 00:01:00 | 1.55 |
|  |  | 75 | 00:00:15 | 0.075 |

**Supplementary Table 3.** Effects of a range of glyphosate concentrations on the human gut microbiota during a short-term colonic incubation of 48h. Measurements assessed change in pH, change in gas pressure, total SCFA, acetate, propionate, butyrate, branched SCFA and lactate production during the 0-48h time interval upon treatment with different concentrations of glyphosate. Enterobacteriaceae concentrations (16S RNA gene copies/mL) after 0h and 48h of incubation upon treatment with different concentrations of glyphosate are also measured.

| Glyphosate (mg/L) | 0 | 0.5 | 1 | 3 | 10 | 100 | 1000 |
| --- | --- | --- | --- | --- | --- | --- | --- |
| pH change | -0.47 | -0.49 | -0.52 | -0.35 | -0.46 | -0.54 | -0.32 |
| Gas (pKa) | 116.6 | 111.2 | 110.9 | 112.4 | 112.4 | 111.5 | 130.1 |
| Total SCFA (mM) | 44.0 | 44.1 | 43.8 | 44.1 | 44.4 | 42.8 | 41.2 |
| Acetate (mM) | 16.5 | 17.1 | 16.7 | 16.7 | 17.1 | 16.2 | 14.3 |
| Propionate (mM) | 13.7 | 13.7 | 13.6 | 13.7 | 13.9 | 13.4 | 12.0 |
| Butyrate (mM) | 12.9 | 12.7 | 12.6 | 12.4 | 12.3 | 12.3 | 14.0 |
| Branched SCFA (mM) | 0.29 | 0.36 | 0.31 | 0.34 | 0.40 | 0.33 | 0.20 |
| Lactate (mM) | 0.65 | 0.71 | 0.79 | 0.54 | 0.65 | 0.38 | 0.56 |
| Enterobacteriaceae 0h | 1.3x10<sup>8</sup> | 8.8x10<sup>7</sup> | 1.1x10<sup>8</sup> | 1.0x10<sup>8</sup> | 1.0x10<sup>8</sup> | 1.4x10<sup>8</sup> | 1.5x10<sup>8</sup> |
| Enterobacteriaceae 48h | 2.0x10<sup>10</sup> | 2.4x10<sup>10</sup> | 1.9x10<sup>10</sup> | 2.3x10<sup>10</sup> | 2.1x10<sup>10</sup> | 2.2x10<sup>10</sup> | 1.8x10<sup>10</sup> |

**Supplementary Table S4.** *Firmicutes/Bacteroidetes ratio in ascending, transverse and descending colon compartments during the different experimental periods. Values represent the average ± standard deviation (control n = 6; treatment n = 9; co-treatment n = 9). Statistically significant differences are marked in bold*.

|  |  | Firmicutes/Bacteroidetes ratio |  |  |  |  |  |  |  |  |  |  |  |  |  |  |  |  |  |
| --- | --- | --- | --- | --- | --- | --- | --- | --- | --- | --- | --- | --- | --- | --- | --- | --- | --- | --- | --- |
|  |  | Lumen |  |  |  |  |  |  |  |  | Mucus |  |  |  |  |  |  |  |  |
|  |  | Control |  |  | Treatment |  |  | Co-treatment |  |  | Control |  |  | Treatment |  |  | Co-treatment |  |  |
| AC | Glyphosate | 0,35 | ± | 0,03 | **0,25** | ± | **0,07** | **0,23** | ± | **0,03** | 1,33 | ± | 0,45 | **0,98** | ± | **0,22** | **0,77** | ± | **0,13** |
|  | Roundup | 0,24 | ± | 0,16 | 63,13 | ± | 272,12 | **1,42** | ± | **0,32** | 0,72 | ± | 0,39 | **14,95** | ± | **7,05** | **4,87** | ± | **0,71** |
| TC | Glyphosate | 0,52 | ± | 0,25 | 0,45 | ± | 0,32 | 0,42 | ± | 0,09 | 3,42 | ± | 0,25 | **1,86** | ± | **0,55** | **1,45** | ± | **0,21** |
|  | Roundup | 0,45 | ± | 0,07 | 3,36 | ± | 4,07 | **0,79** | ± | **0,24** | 2,76 | ± | 0,34 | **3,98** | ± | **1,27** | **3,48** | ± | **0,52** |
| DC | Glyphosate | 0,87 | ± | 0,04 | **0,69** | ± | **0,05** | **0,44** | ± | **0,10** | 3,47 | ± | 1,51 | **1,77** | ± | **0,19** | **1,27** | ± | **0,21** |
|  | Roundup | 0,71 | ± | 0,19 | **3,36** | ± | **4,07** | **1,12** | ± | **0,21** | 3,44 | ± | 0,86 | 3,98 | ± | 1,27 | **4,30** | ± | **0,21** |

