## Supplementary Figures S8 for "Alterations in human gut microbiome composition and metabolism after exposure to glyphosate and Roundup and/or a spore-based formulation using the SHIME® technology"

### Metabolome plots

```
## [1] "X-25656"
```

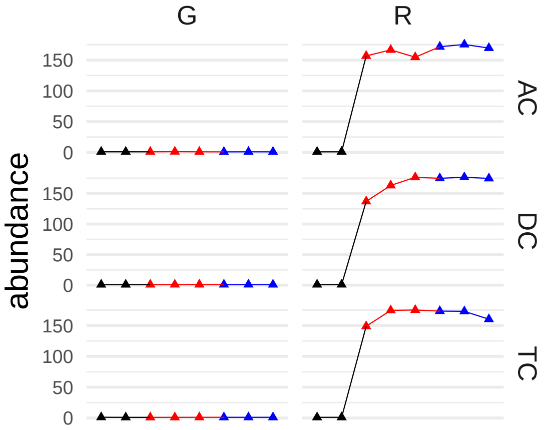


```
## [1] "sphingosine"
```

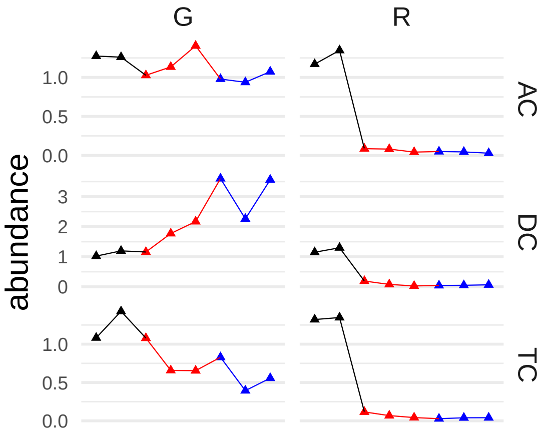


```
## [1] "glyphosate"
```

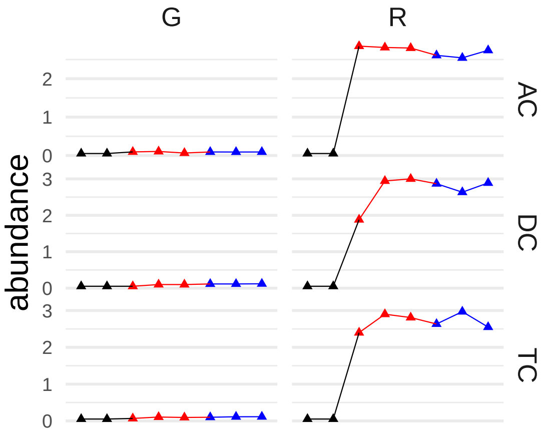

```
## [1] "glycerol 3-phosphate"
```

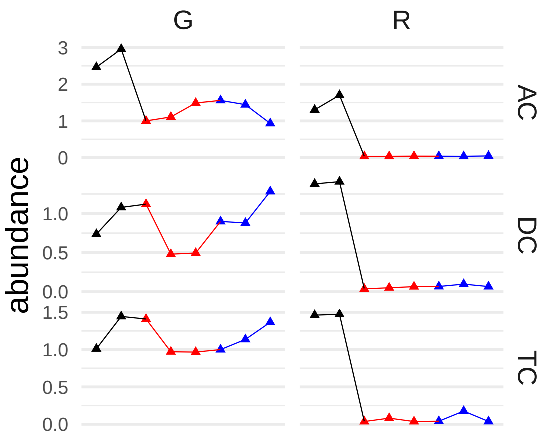


```
## [1] "ethylmalonate"
```

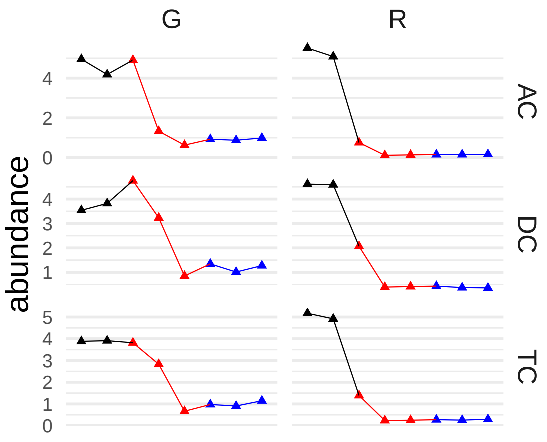


```
## [1] "2-pyrrolidinone"
```

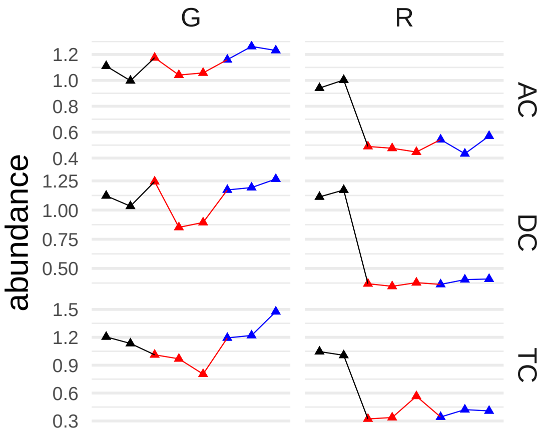


```
## [1] "cyclo(phe-pro)"
```

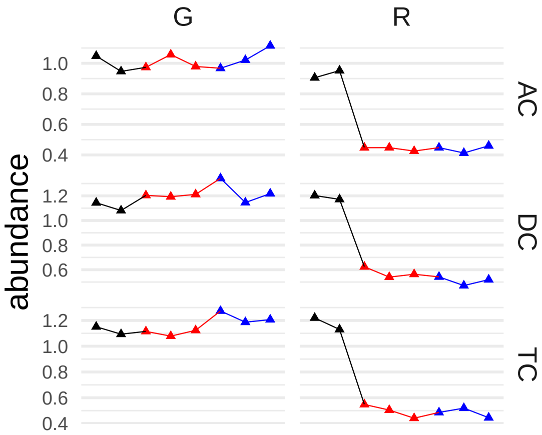


#### [1] "sphinganine"

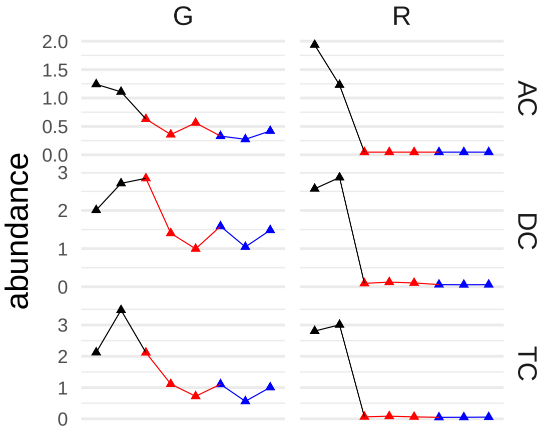


#### [1] "valerate (5:0)"

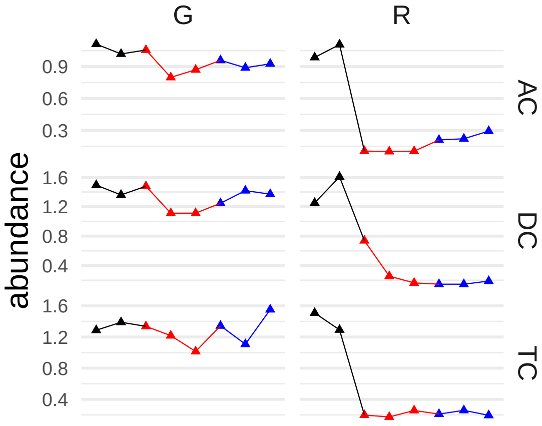


#### [1] "methylphosphate"

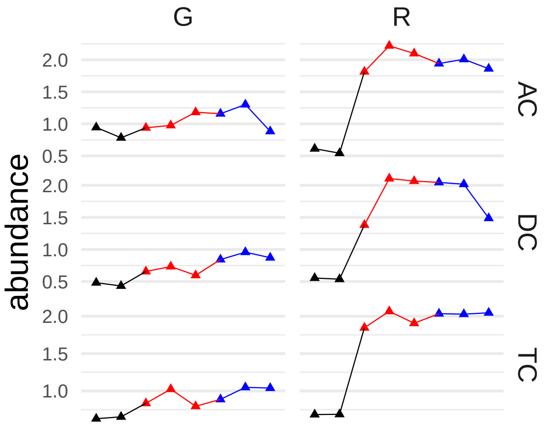


```
## [1] "indoleacetate"
```

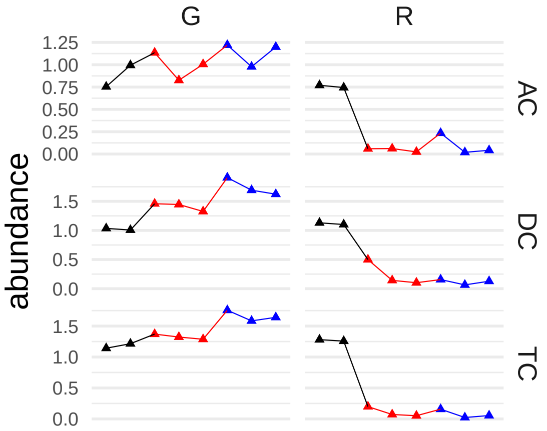


```
## [1] "butyrate/isobutyrate (4:0)"
```

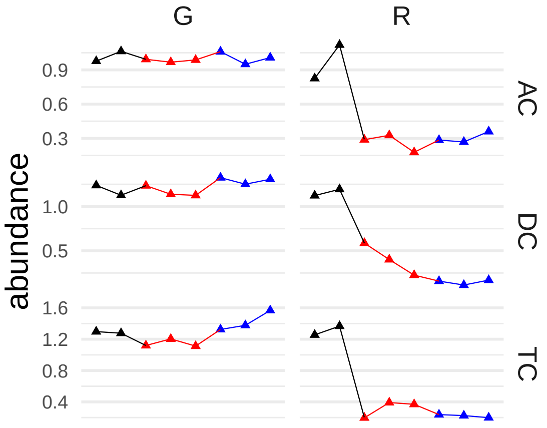


```
## [1] "ribose"
```

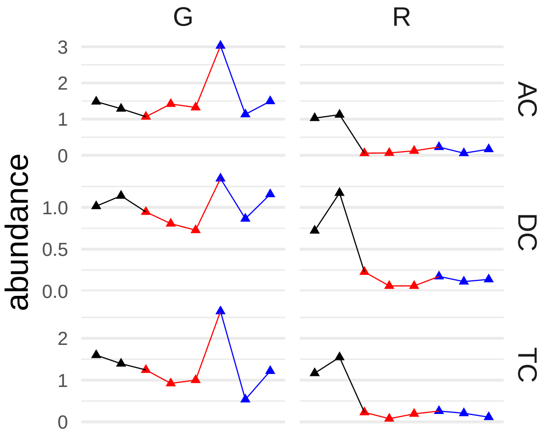


#### [1] "3-ketosphinganine"

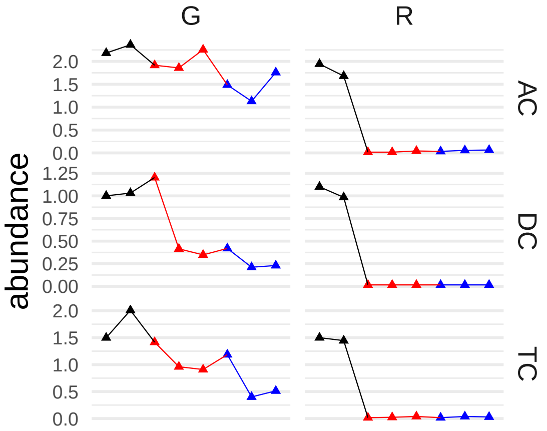


#### [1] "sphingomyelin (d17:1/16:0, d18:1/15:0, d16:1/17:0)*"

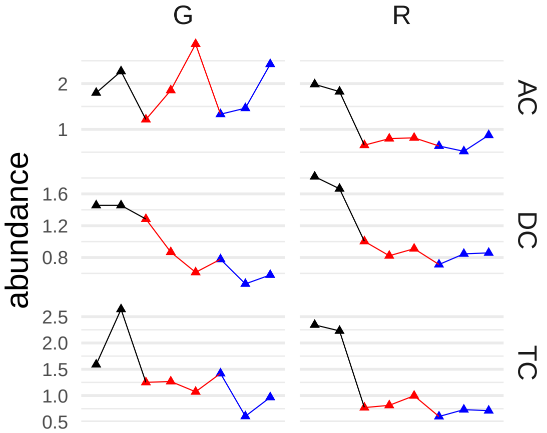


#### [1] "1,3-propanediol"

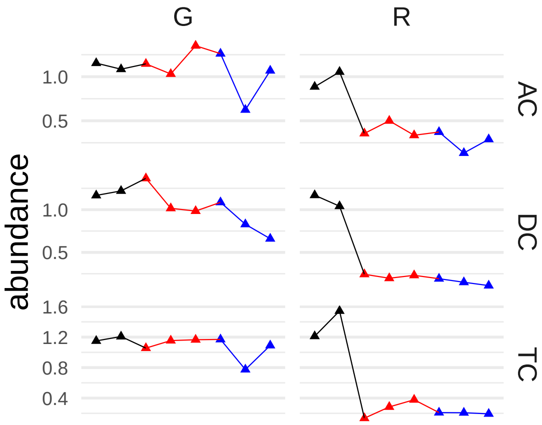


```
## [1] "cyclo(leu-pro)"
```

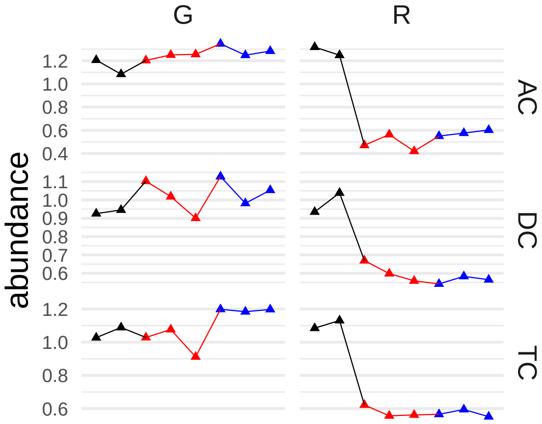


```
## [1] "methylsuccinate"
```

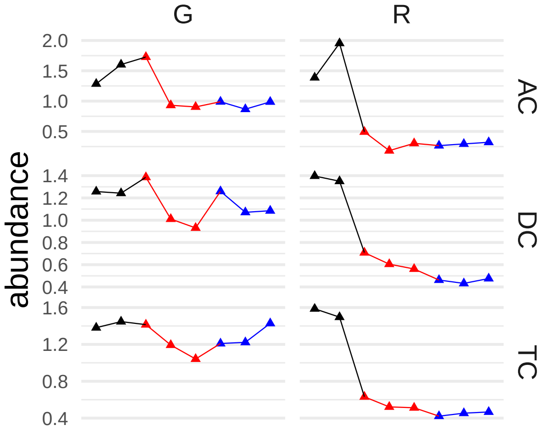


```
## [1] "oleoyl ethanolamide"
```

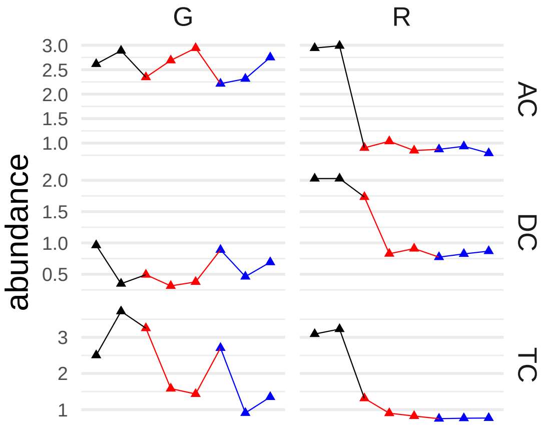


```
## [1] "4-hydroxycinnamate"
```

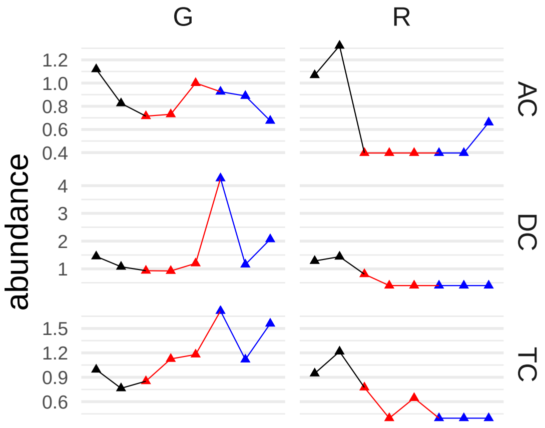


```
## [1] "X-24731"
```

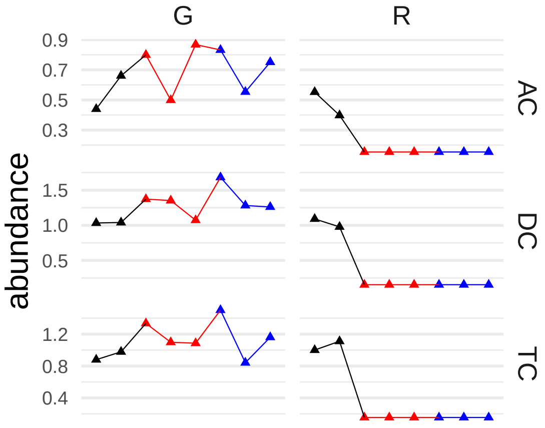


```
## [1] "cyclo(phe-pro) (L,D)*"
```

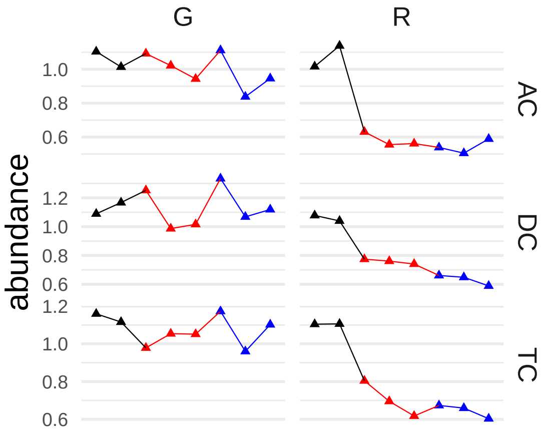


#### [1] "gamma-tocopherol/beta-tocopherol"

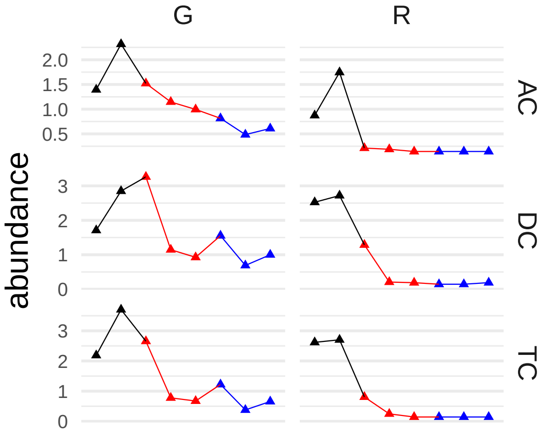


#### [1] "phosphate"

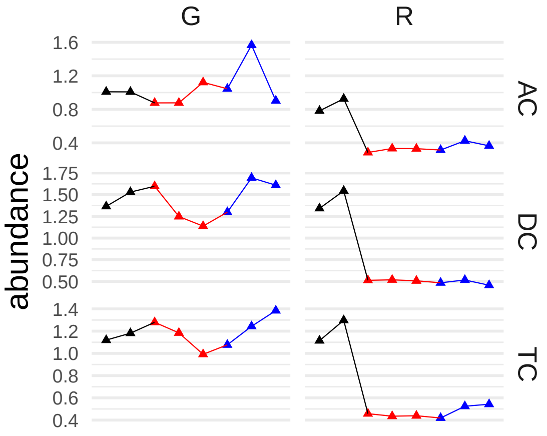


#### [1] "caproate (6:0)"

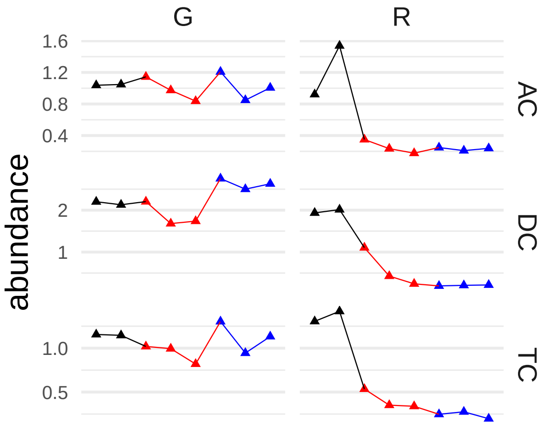


#### [1] "tryptamine"

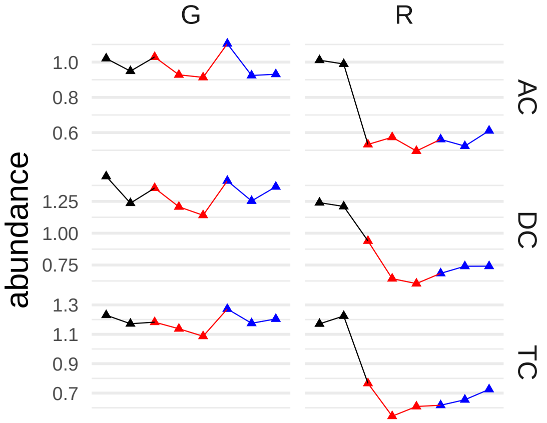


#### [1] "2-oxindole-3-acetate"

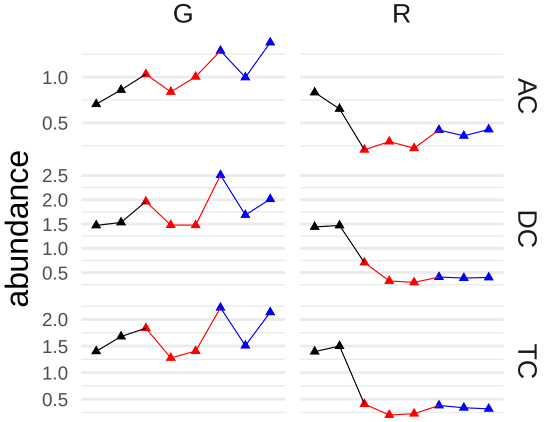


#### [1] "3-hydroxyarachidate*"

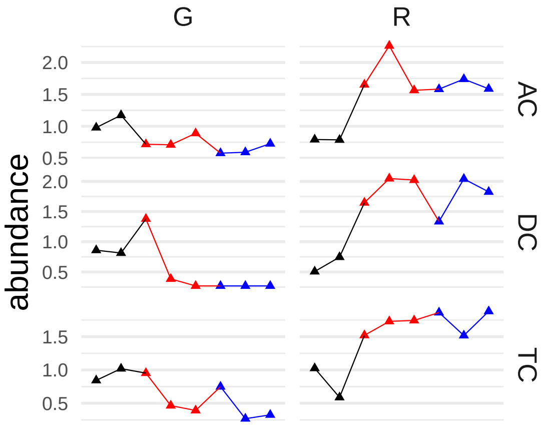


#### [1] "maltol"

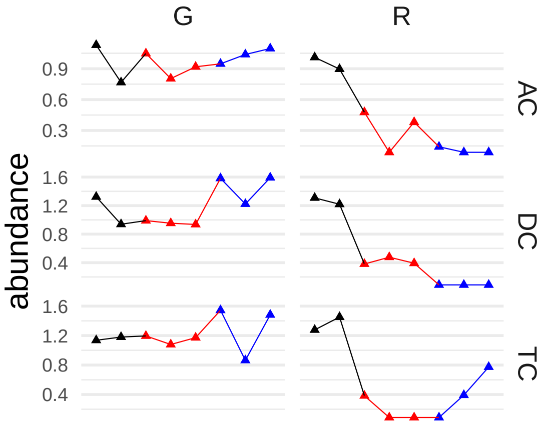


#### [1] "N6,N6-dimethyllysine"

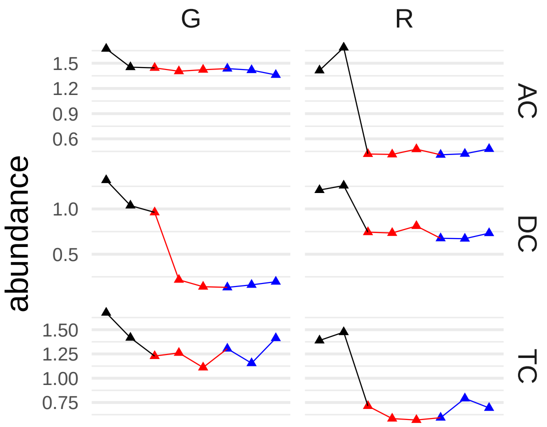


#### [1] "uridine 5'-monophosphate (UMP)"

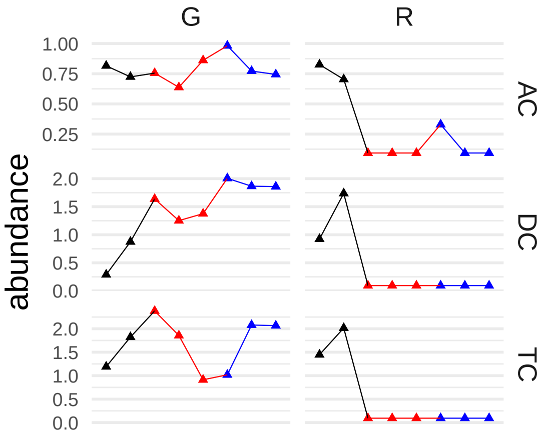


```
## [1] "isovalerate (i5:0)"
```

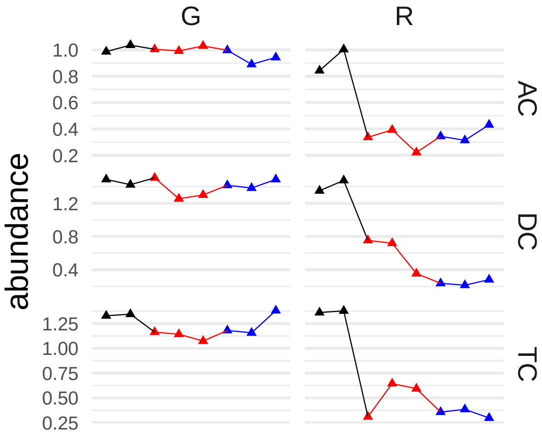


```
## [1] "2,4-di-tert-butylphenol"
```

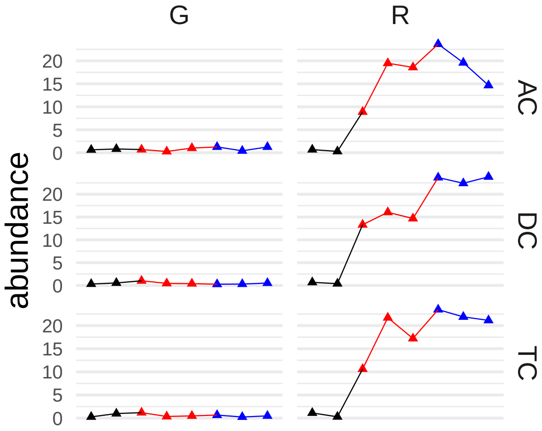


```
## [1] "N-acetylaspartate (NAA)"
```

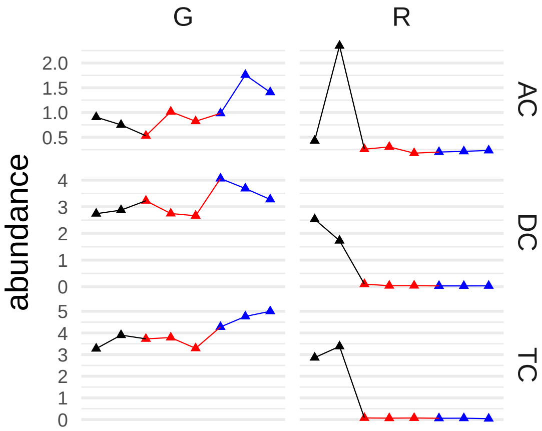


```
## [1] "indole-3-carboxylate"
```

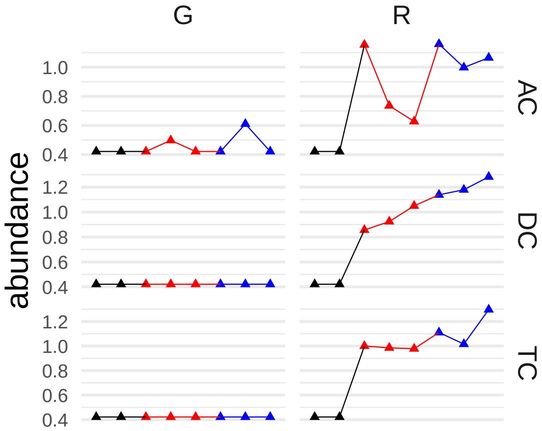


```
## [1] "X-25594"
```

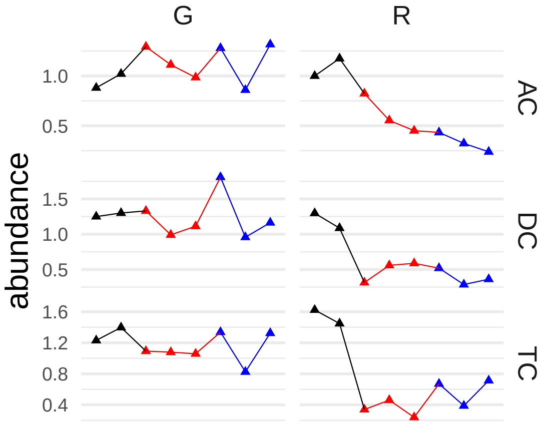


```
## [1] "X-15956"
```

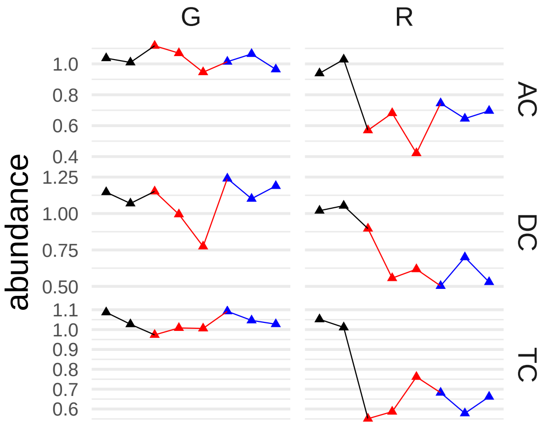


```
## [1] "cholesterol sulfate"
```

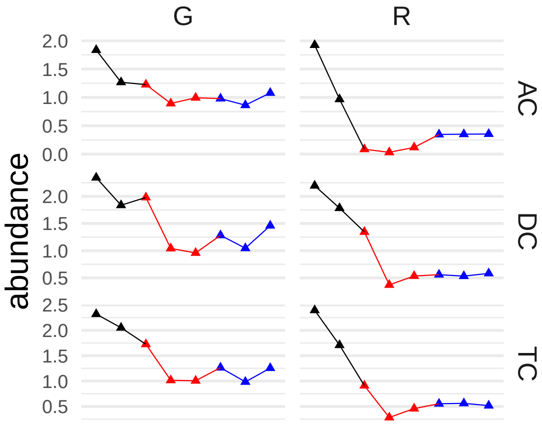


```
## [1] "stearoyl ethanolamide"
```

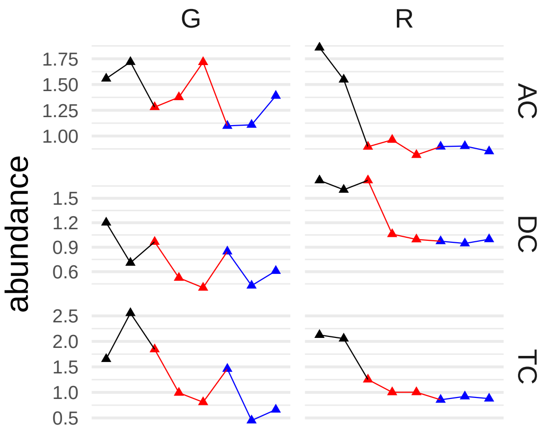


```
## [1] "tyramine"
```

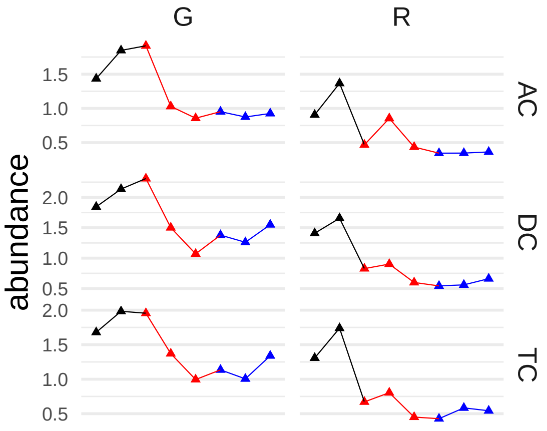


```
## [1] "pentose acid*"
```

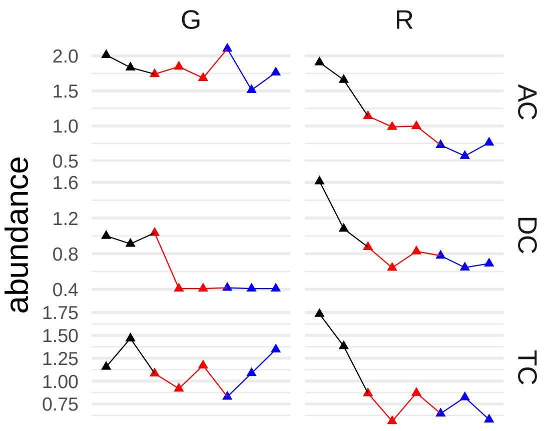


```
## [1] "docosapentaenoate (n3 DPA; 22:5n3)"
```

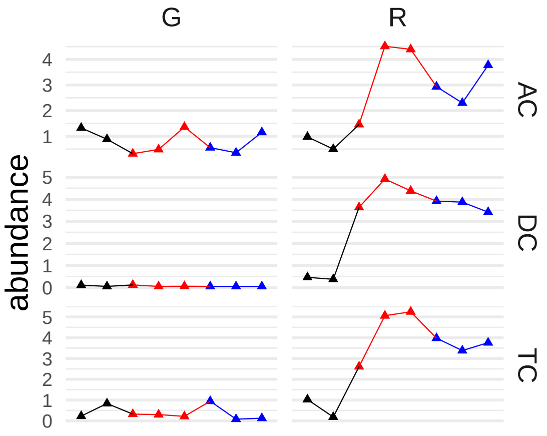


```
## [1] "3-hydroxy-2-methylpyridine"
```

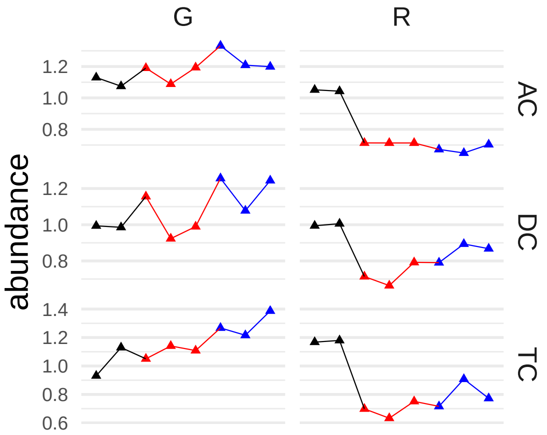


```
## [1] "pyridoxamine"
```

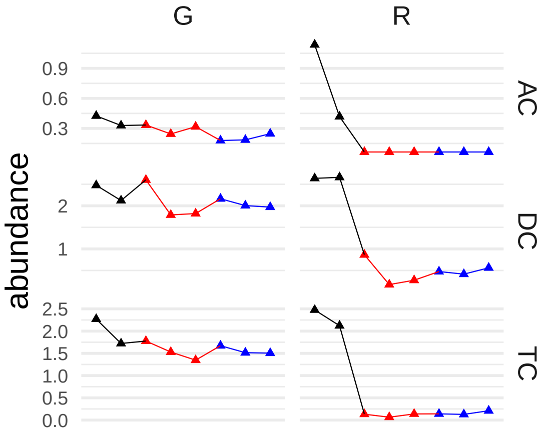


```
## [1] "X-25830"
```

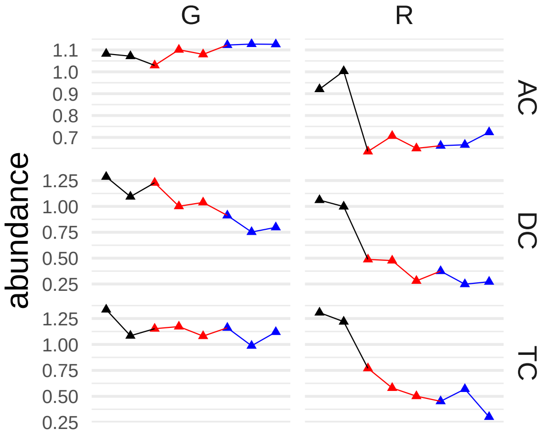


```
## [1] "oleate/vaccenate (18:1)"
```

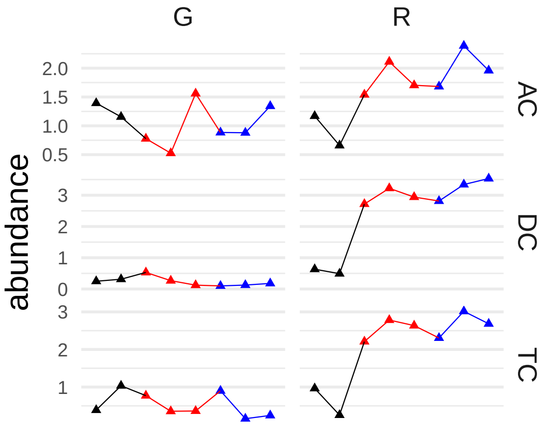


## [1] "X-23732"

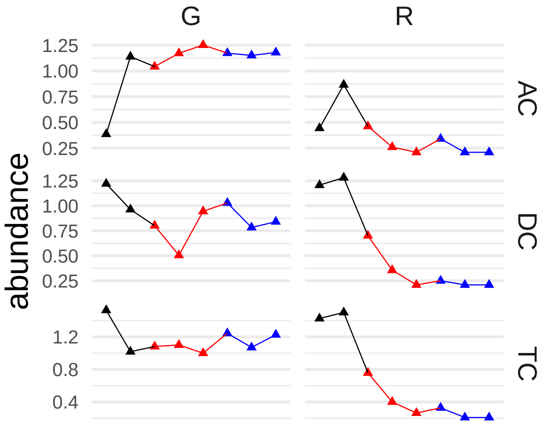


#### [1] "solanidine"

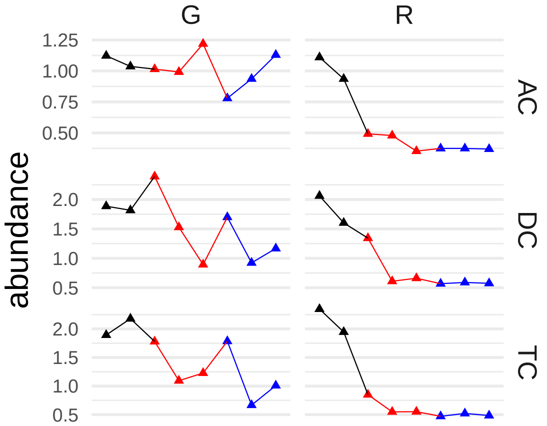


#### [1] "carboxyethyl-GABA"

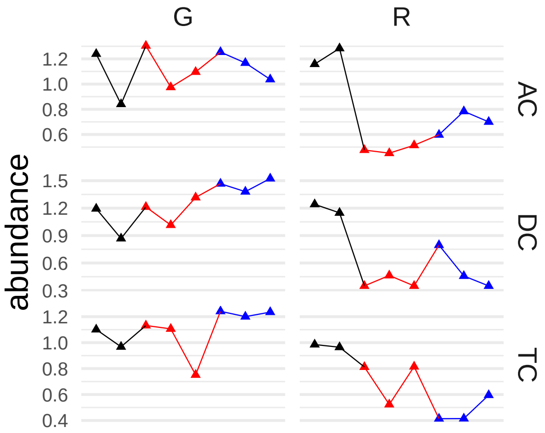


#### [1] "3-hydroxystearate"

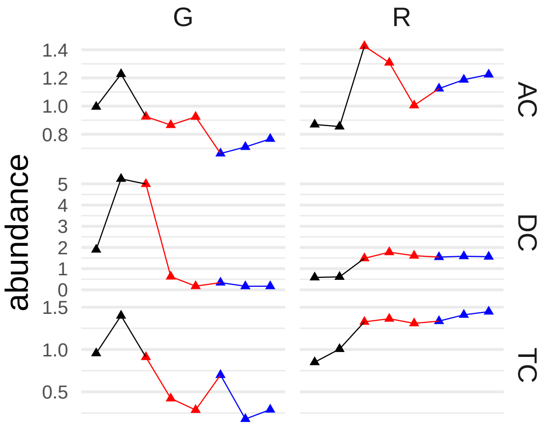


#### [1] "4-ureidobutyrate"

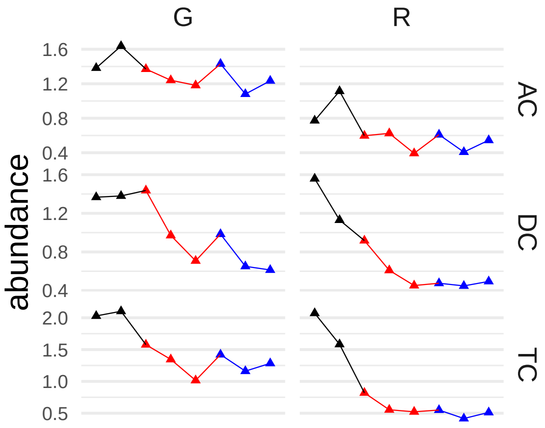


#### [1] "hypoxanthine"

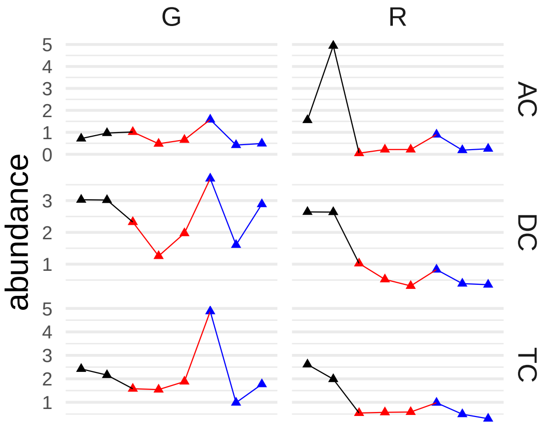


```
## [1] "adenine"
```

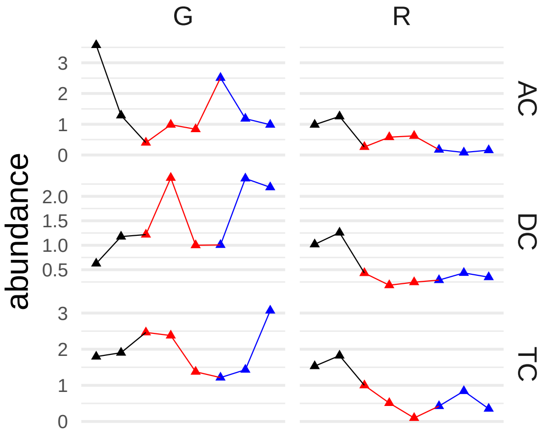


```
## [1] "N-acetyl-cadaverine"
```

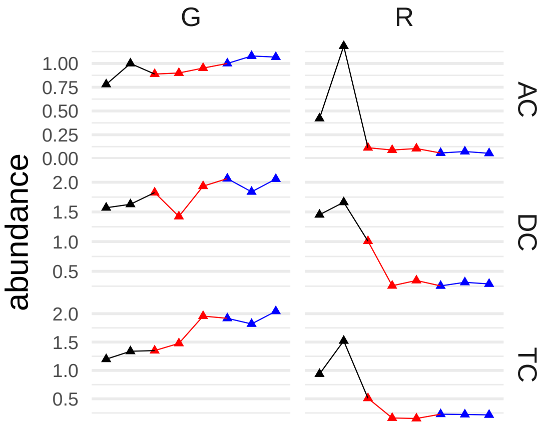


```
## [1] "oxalate (ethanedioate)"
```

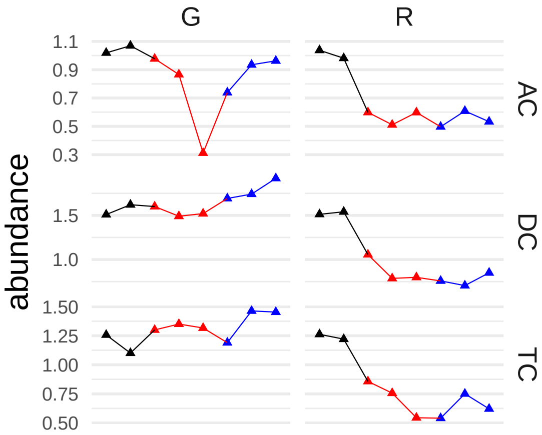


#### [1] "N-carbamoylvaline"

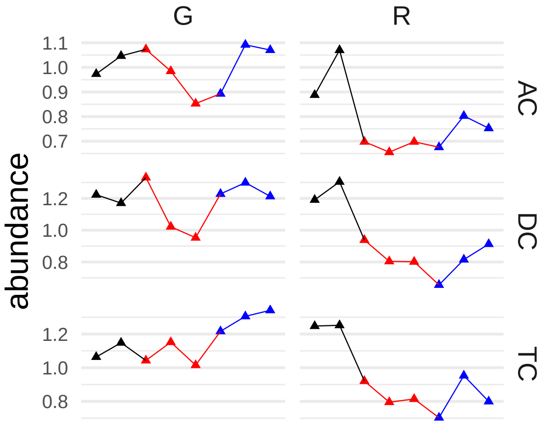


#### [1] "4-guanidinobutanoate"

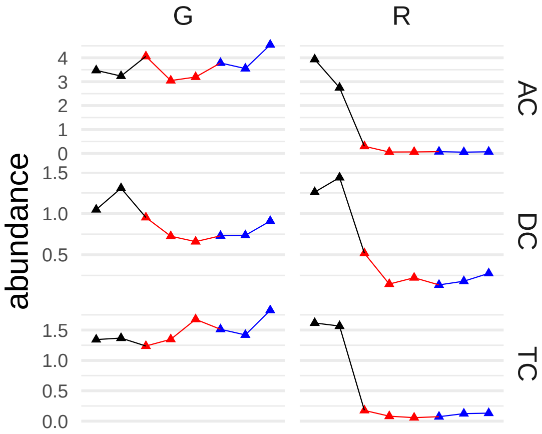


#### [1] "linoleate (18:2n6)"

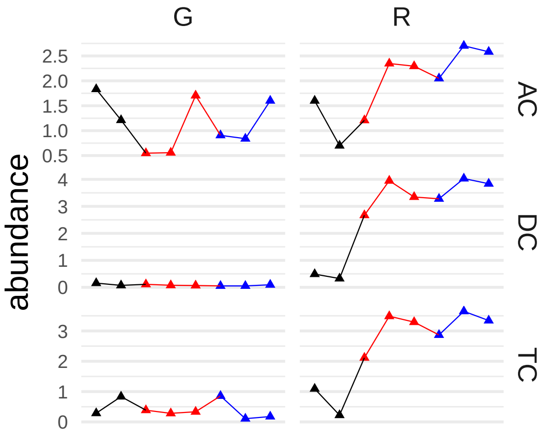


#### [1] "docosahexaenoate (DHA; 22:6n3)"

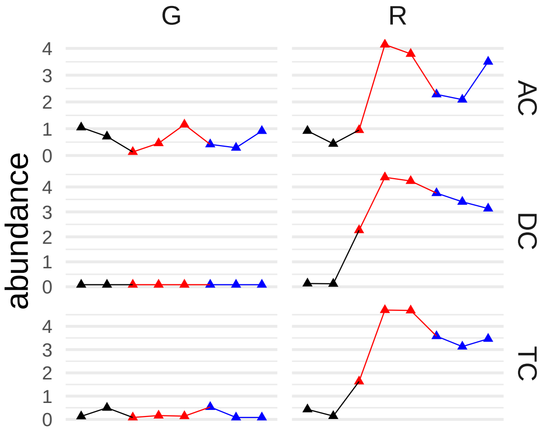


#### [1] "2-iminopiperidine"

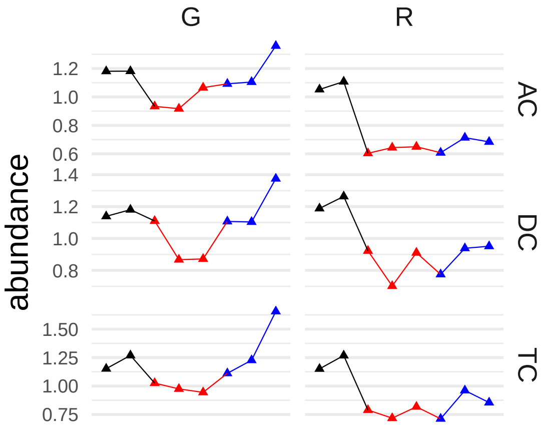


#### [1] "xanthine"

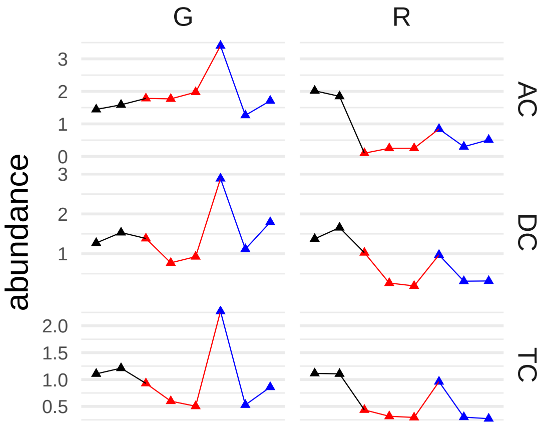


#### [1] "N-acetylmuramate"

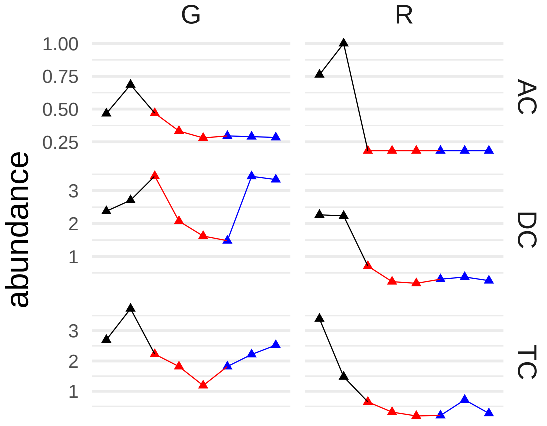


#### [1] "serotonin"

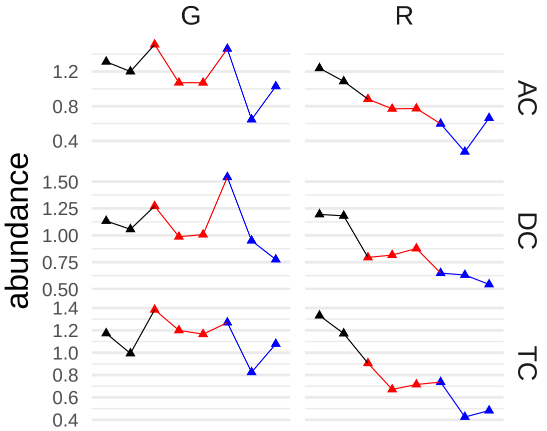


#### [1] "iminodiacetate (IDA)"

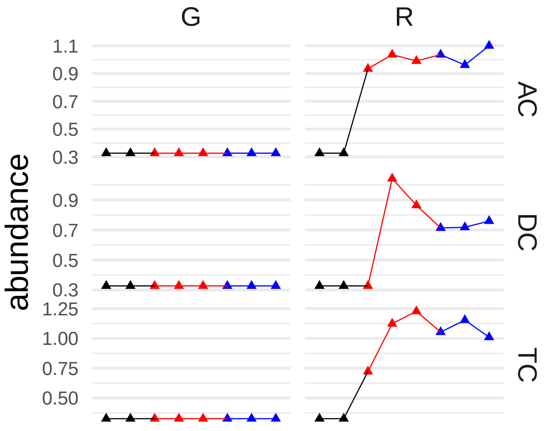


#### [1] "3-formylindole"

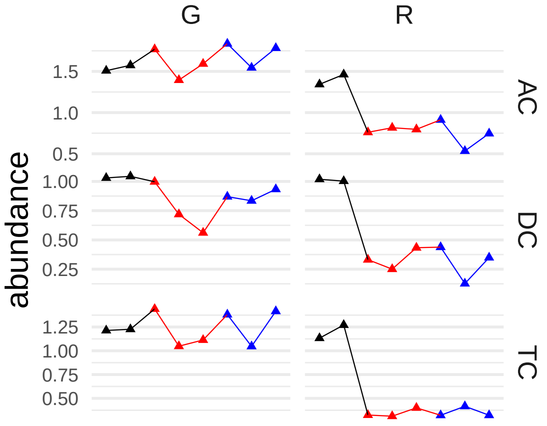


#### [1] "N-acetylputrescine"

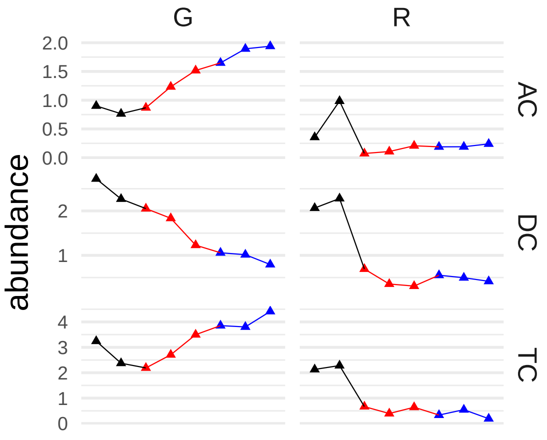


## [1] "X-23662"

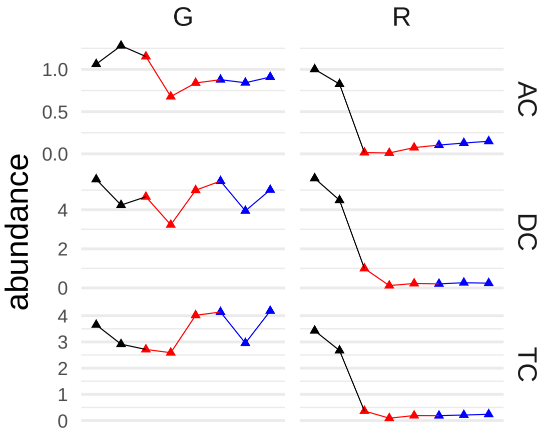


```
## [1] "X-11787"
```

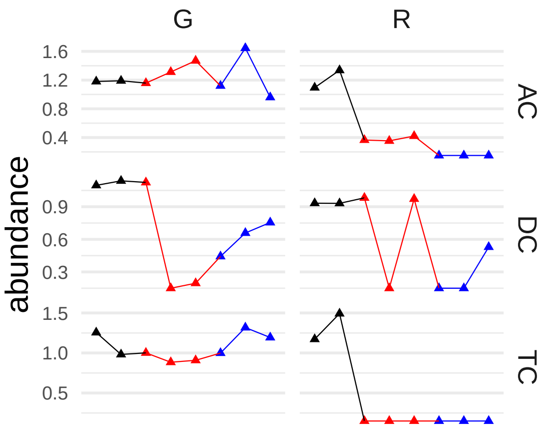


```
## [1] "N-acetyl-1-methylhistidine*"
```

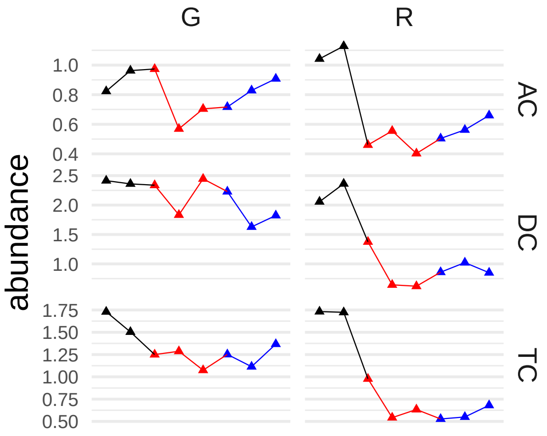


```
## [1] "3-hydroxymyristate"
```

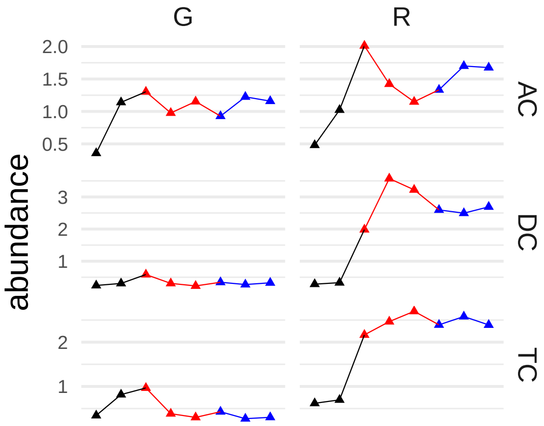


```
## [1] "3-amino-2-piperidone"
```

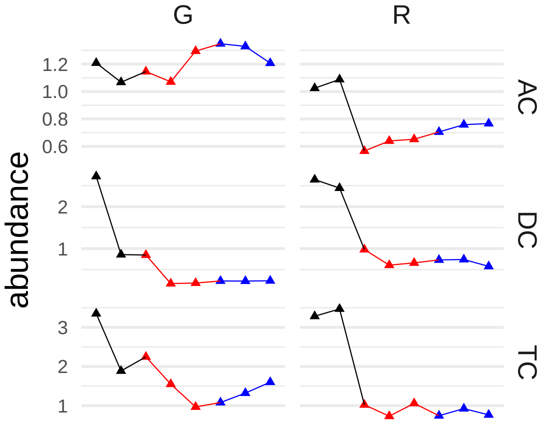


```
## [1] "phenylalanine"
```

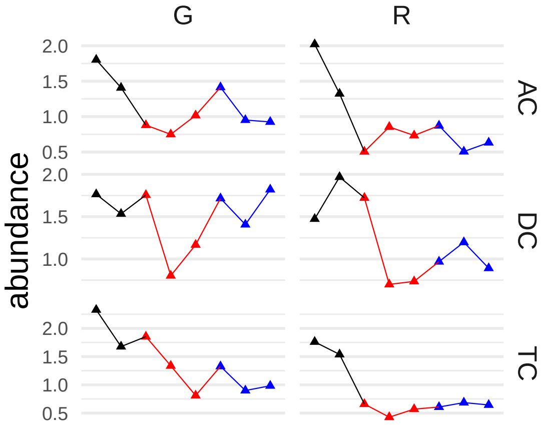


```
## [1] "tyrosine"
```

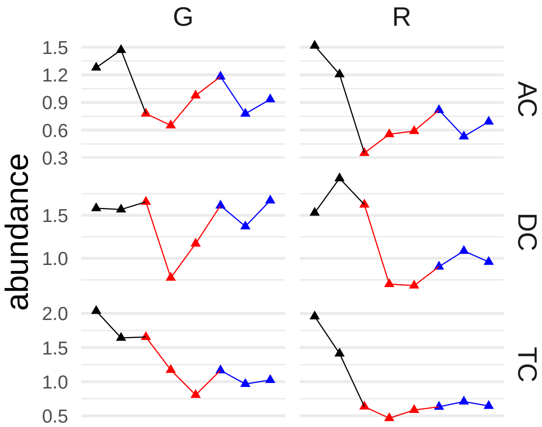


```
## [1] "(S)-a-amino-omega-caprolactam"
```

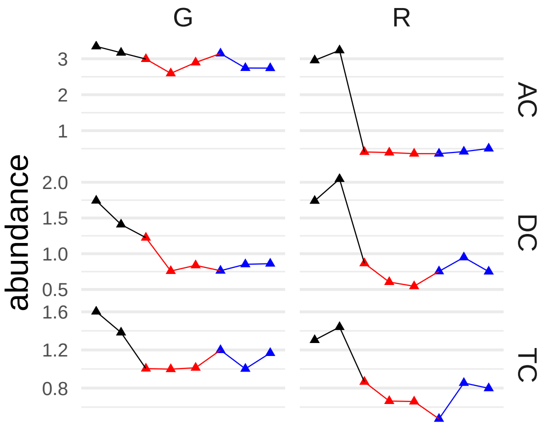


```
## [1] "X-25041"
```

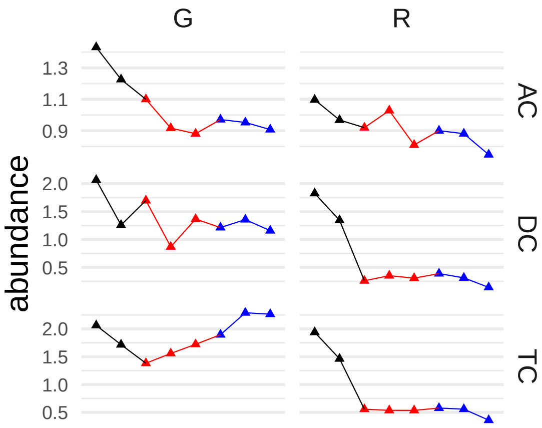


```
## [1] "N-acetylkynurenine (2)"
```

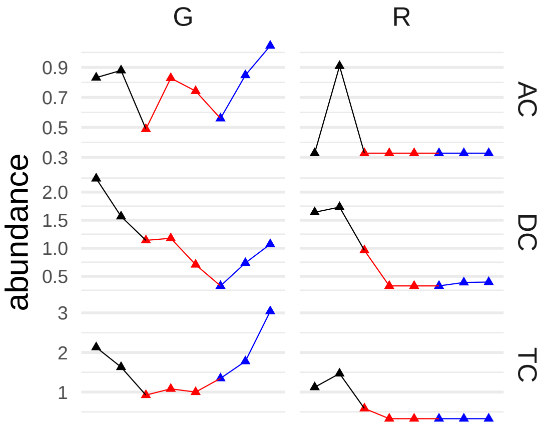


#### [1] "1-methyl-4-imidazoleacetate"

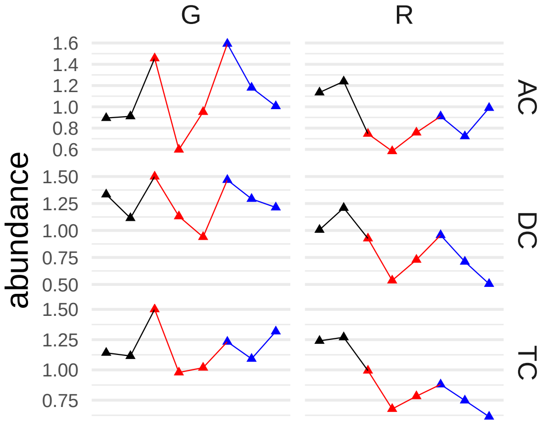


#### [1] "2-heptyl-1H-quinolin-4-one (HHQ)"

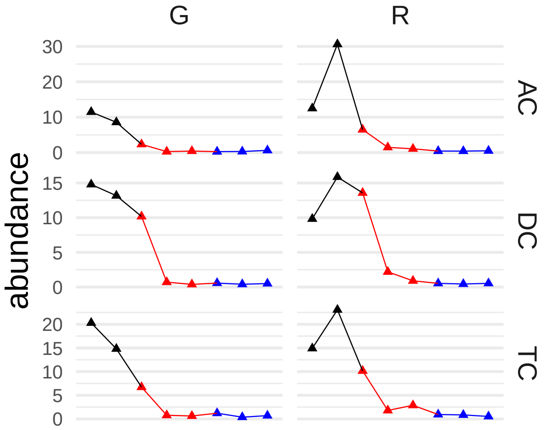


#### [1] "5-methyl-2'-deoxycytidine"

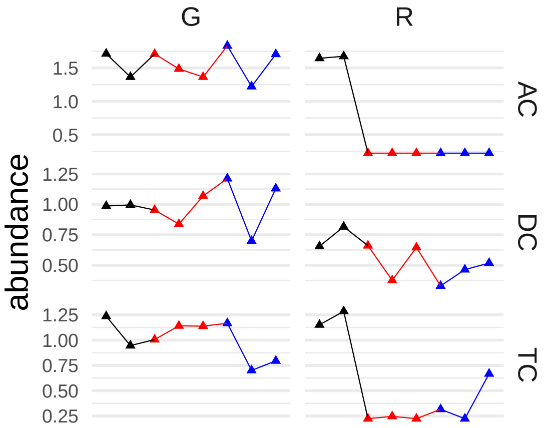


#### [1] "glycosyl-N-palmitoyl-sphingosine (d18:1/16:0)"

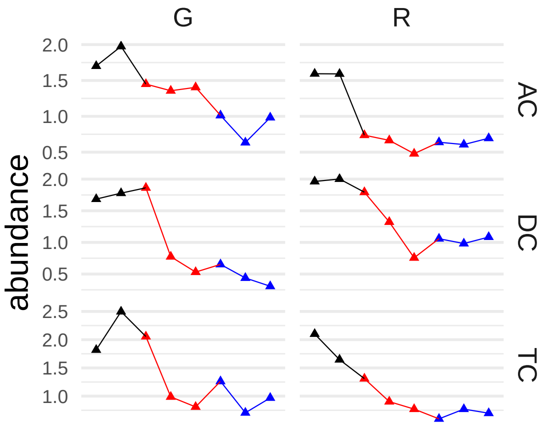


#### [1] "docosapentaenoate (n6 DPA; 22:5n6)"

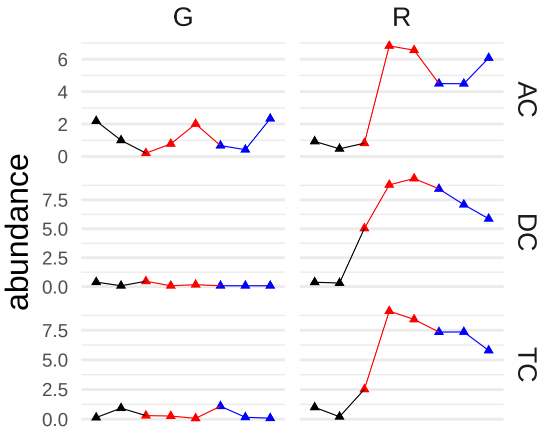


#### [1] "fumarate"

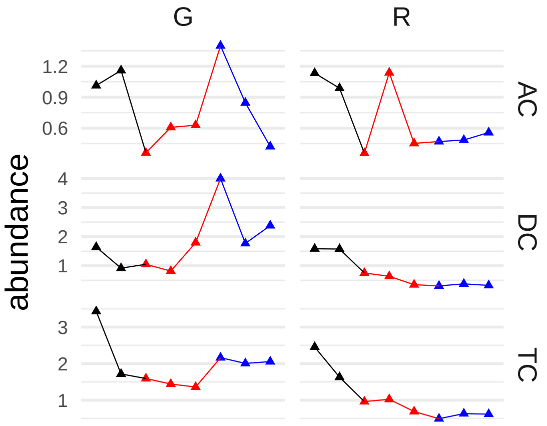


#### [1] "7-hydroxycholesterol (alpha or beta)"

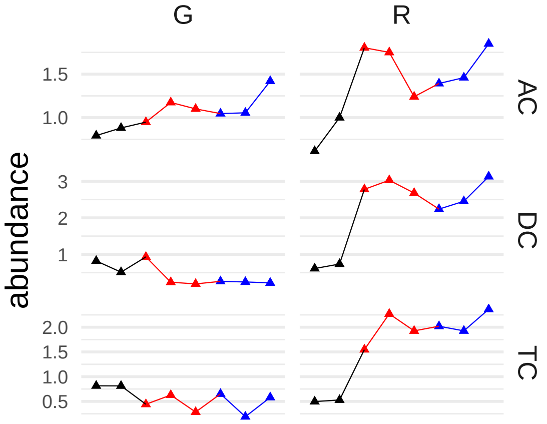


## [1] "X-24237"

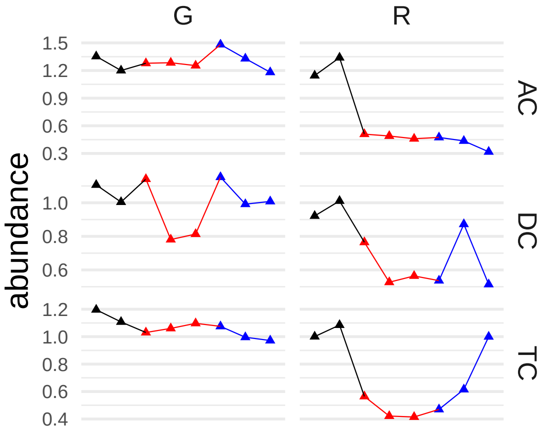


#### [1] "dihomo-linoleate (20:2n6)"

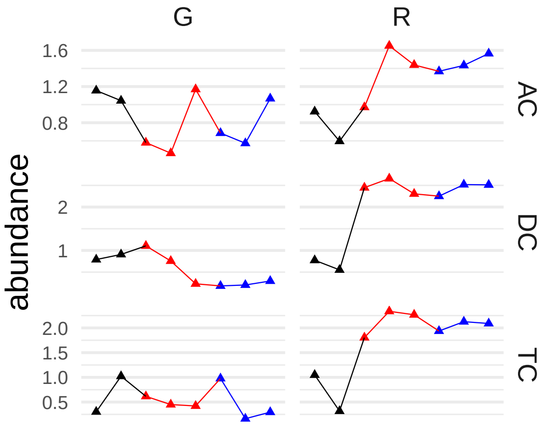


```
## [1] "X-24973"
```

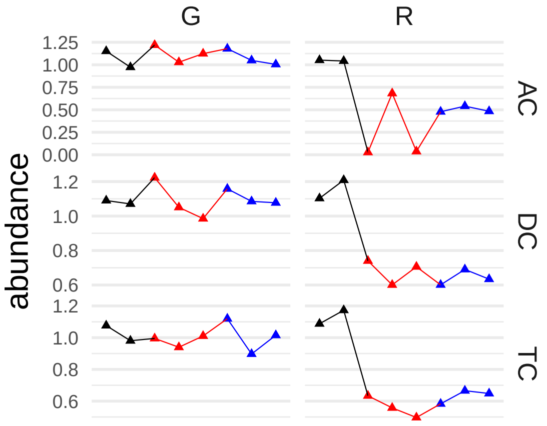


```
## [1] "arachidonate (20:4n6)"
```

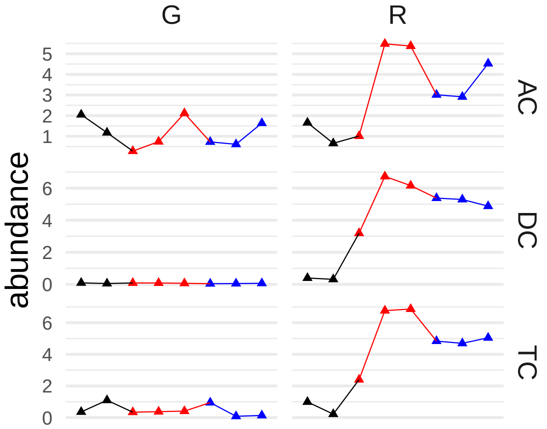


```
## [1] "3-hydroxy-3-methylglutarate"
```

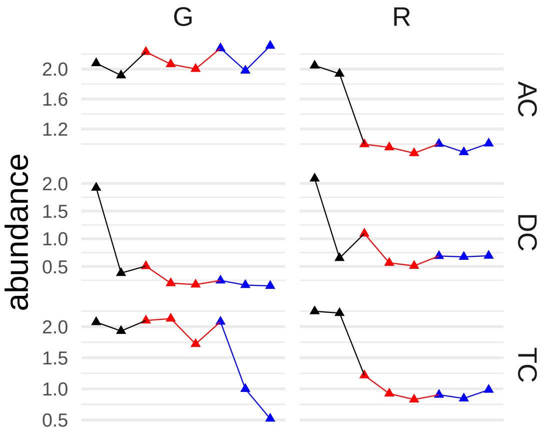


#### [1] "cytidine 5'-monophosphate (5'-CMP)"

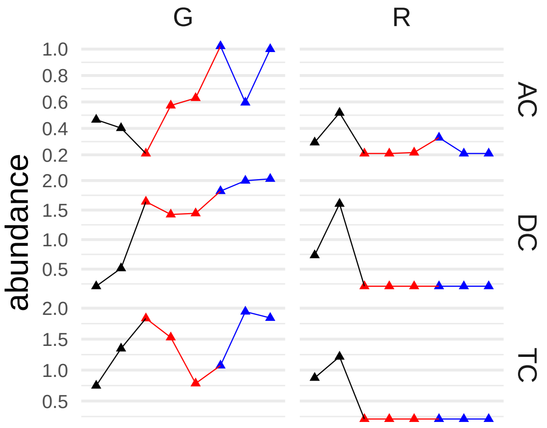


#### [1] "2S,3R-dihydroxybutyrate"

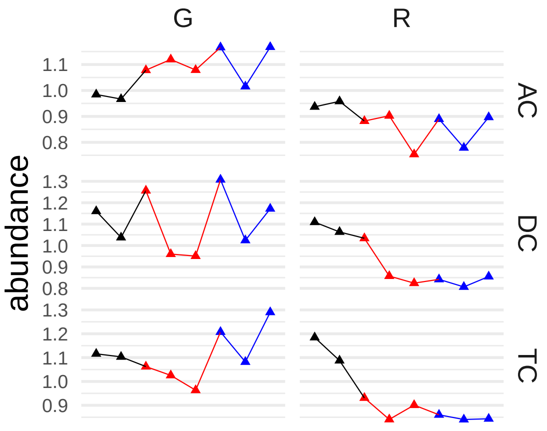


#### [1] "phenethylamine"

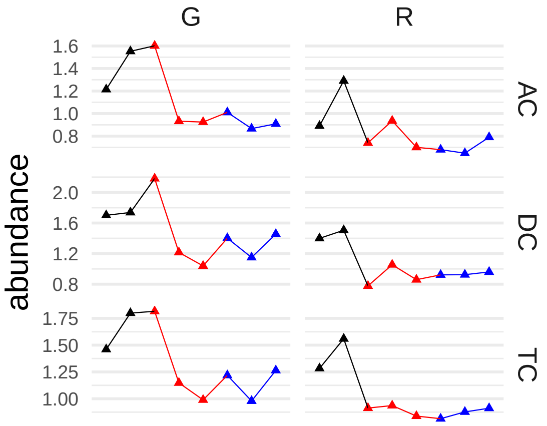


```
## [1] "butyrylputrescine/isobutyrylputrescine"
```

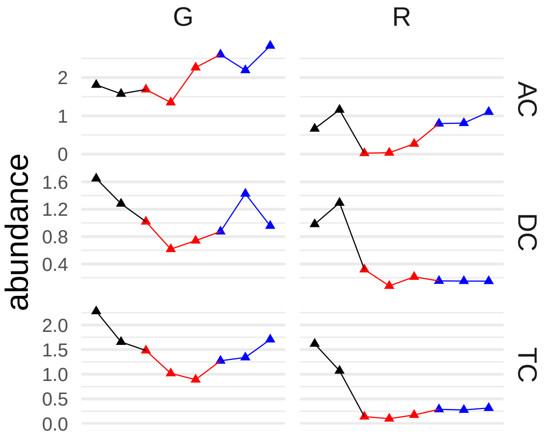


```
## [1] "palmitoyl ethanolamide"
```

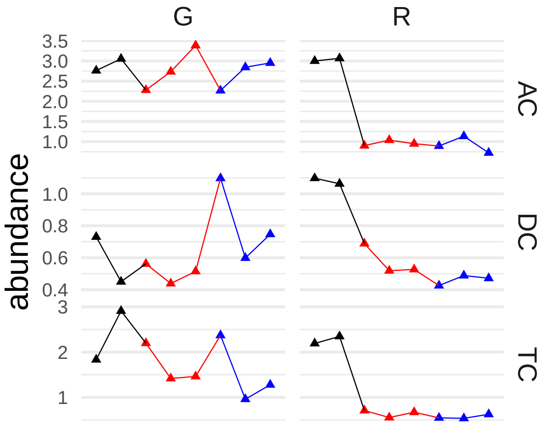


```
## [1] "pyridoxate"
```

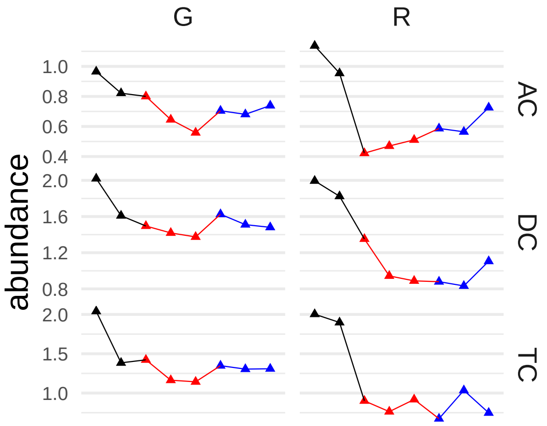


#### [1] "1-methylhypoxanthine"

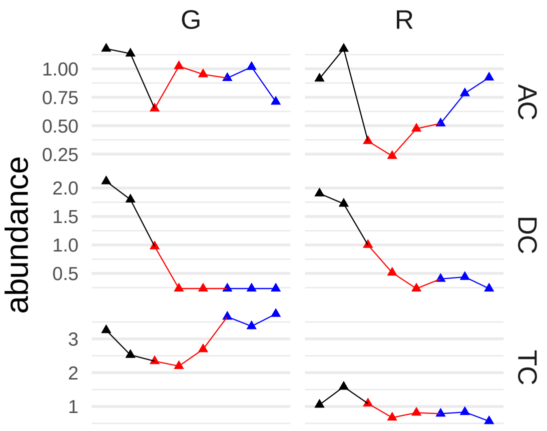


#### [1] "nicotinate ribonucleoside"

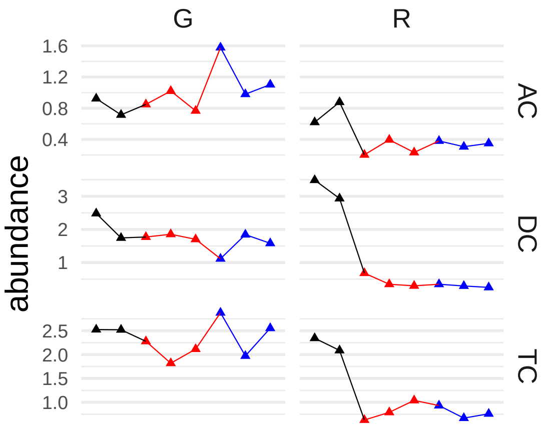


#### [1] "N(1)-acetylspermine"

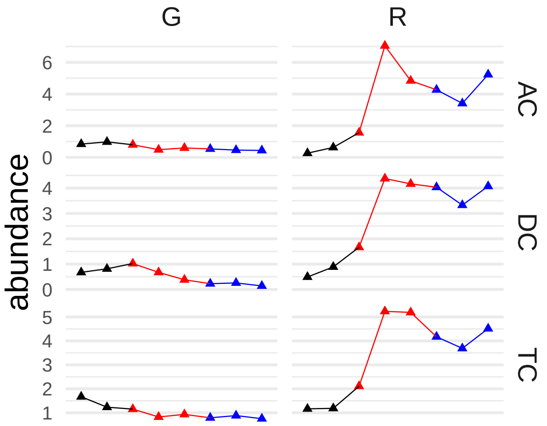


## [1] "X-13507"

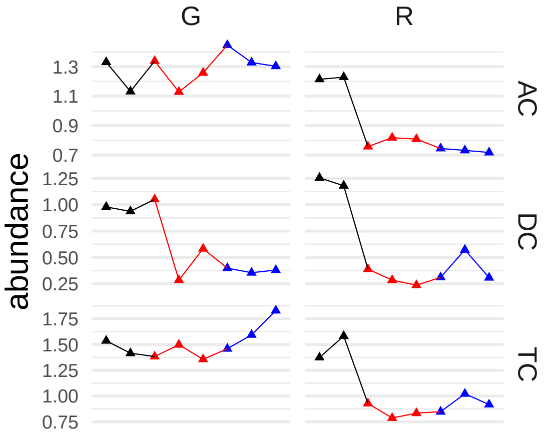


#### [1] "sphingomyelin (d18:2/16:0, d18:1/16:1)*"

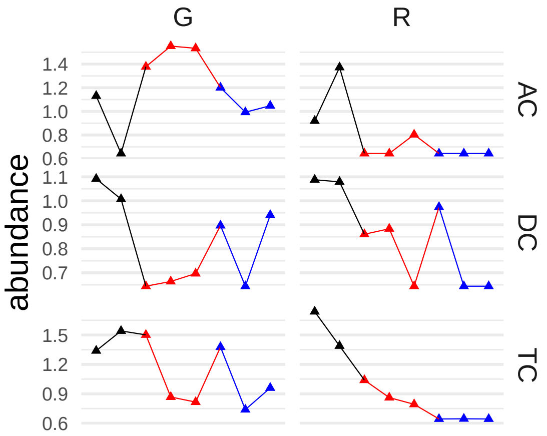


#### [1] "N,N-dimethyl-5-aminovalerate"

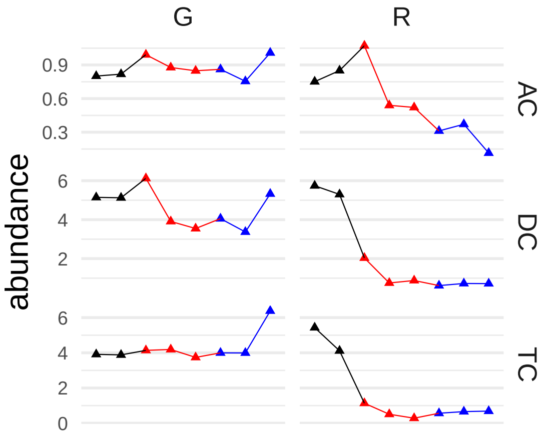


#### [1] "N-acetylhistamine"

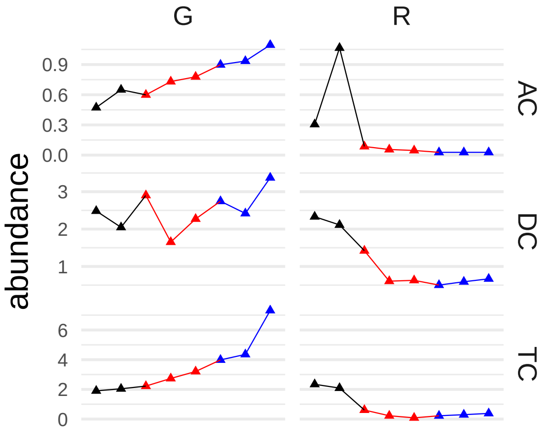


#### [1] "eicosapentaenoate (EPA; 20:5n3)"

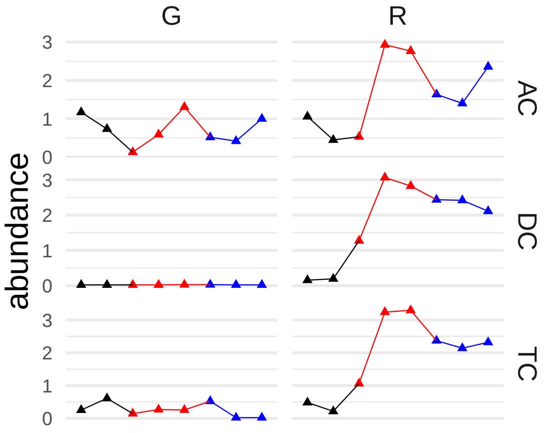


#### [1] "hydantoin-5-propionate"

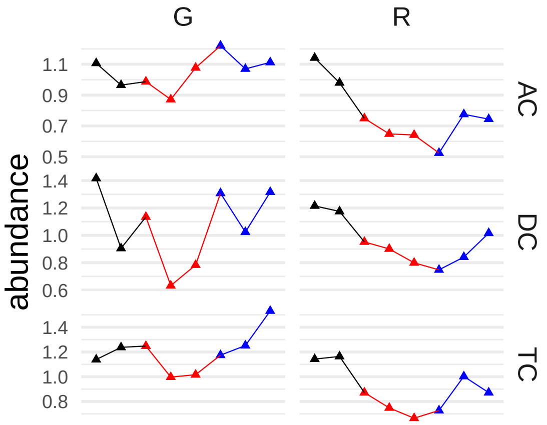


```
## [1] "dihomo-linolenate (20:3n3 or n6)"
```

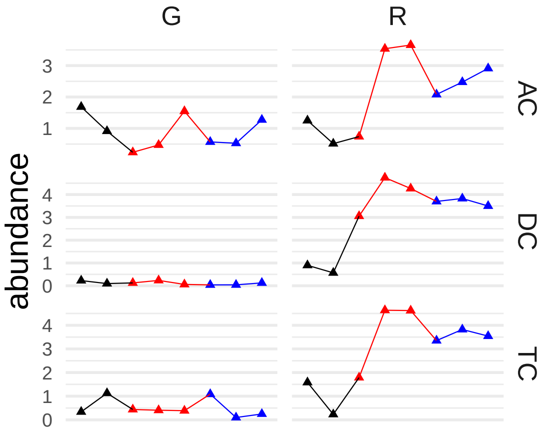


```
## [1] "2'-deoxyadenosine"
```

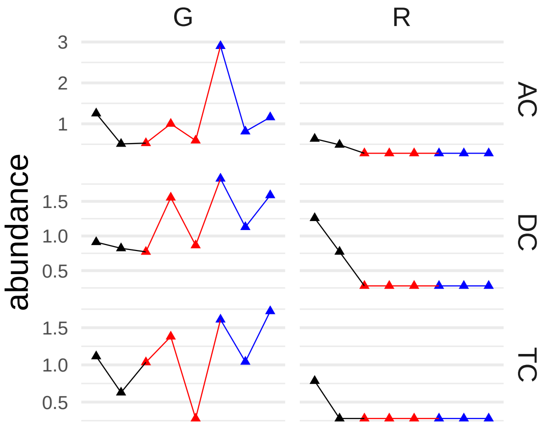


```
## [1] "guanine"
```

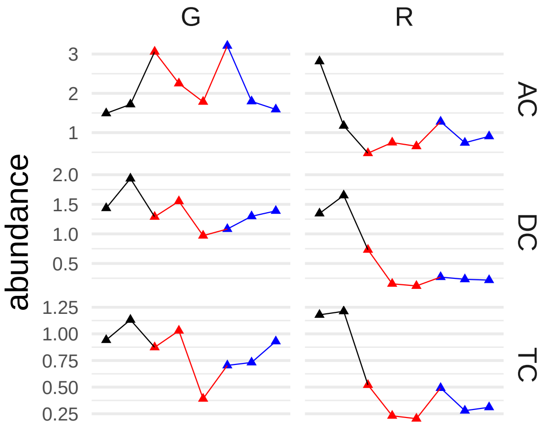


#### [1] "6-oxopiperidine-2-carboxylate"

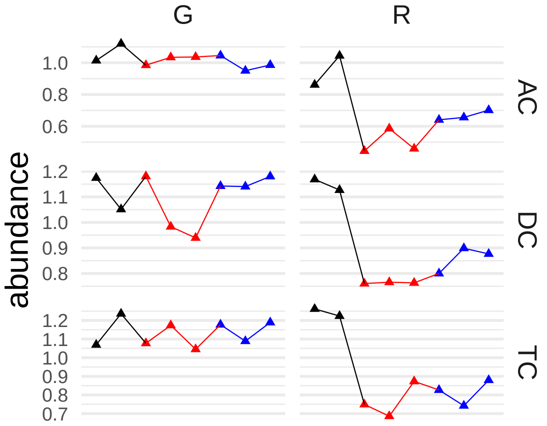


#### [1] "daidzein"

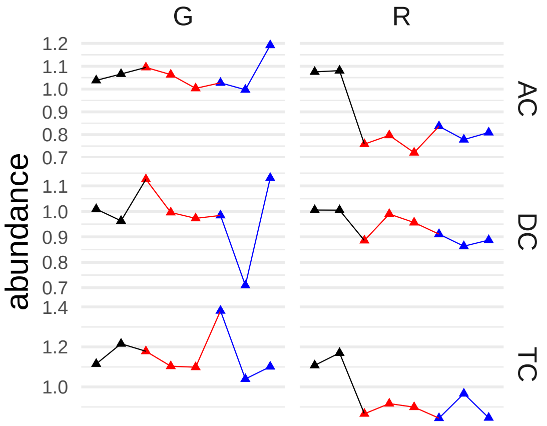


#### [1] "N6-carboxymethyllysine"

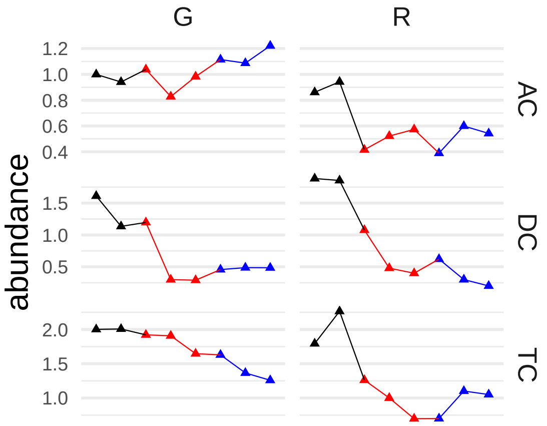


#### [1] "N,N,N-trimethyl-5-aminovalerate"

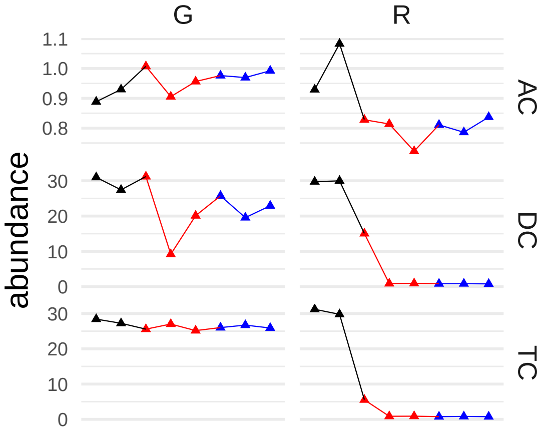


## [1] "X-23240"

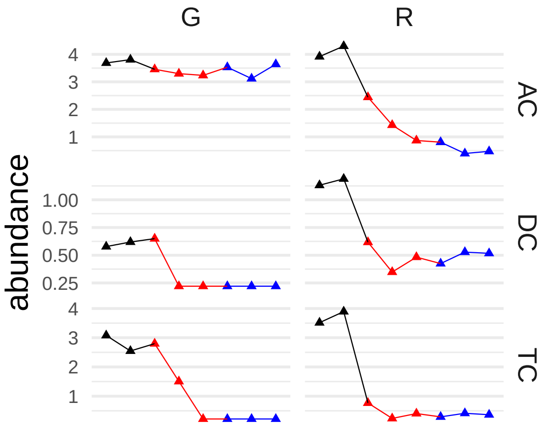


#### [1] "creatine"

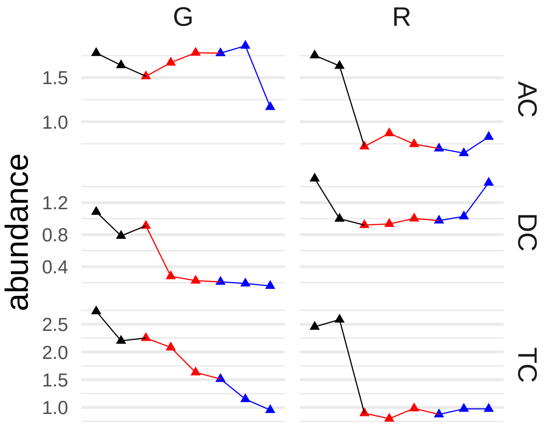


```
## [1] "palmitate (16:0)"
```

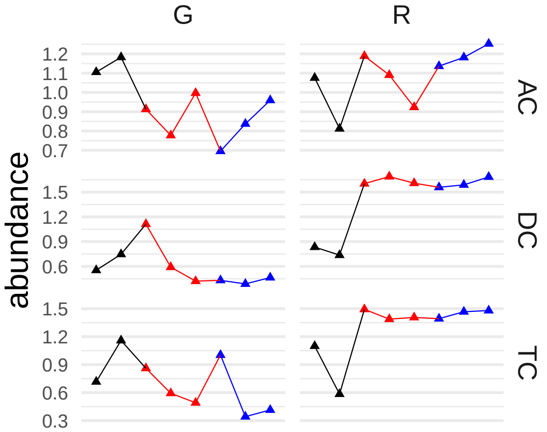


```
## [1] "cis-4-decenoate (10:1n6)*"
```

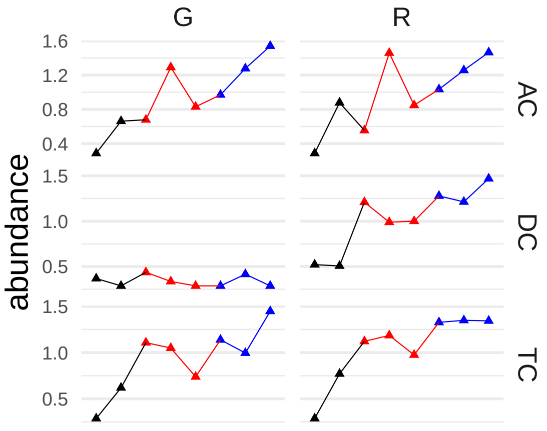


```
## [1] "eicosenoate (20:1)"
```

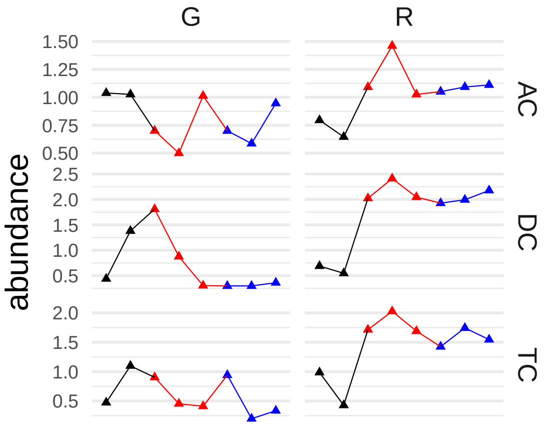


```
## [1] "X-25008"
```

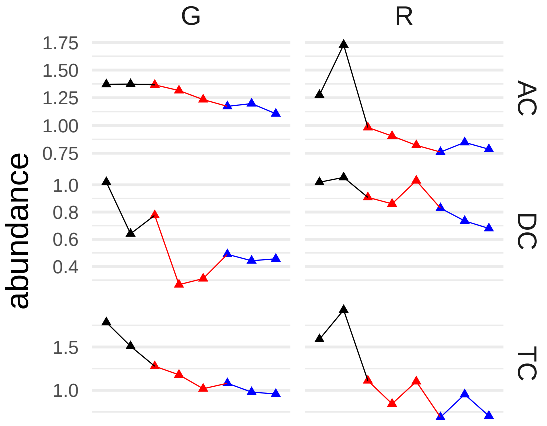


```
## [1] "stearate (18:0)"
```

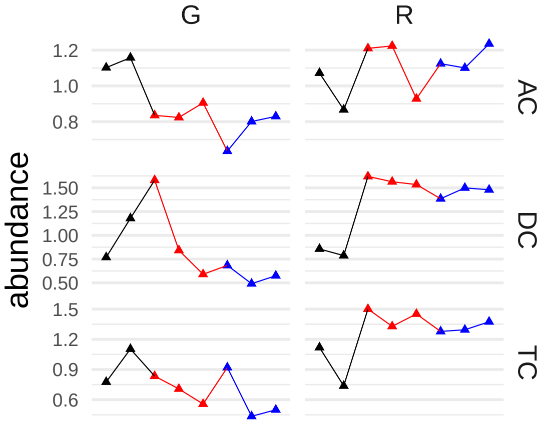


```
## [1] "X-15461"
```

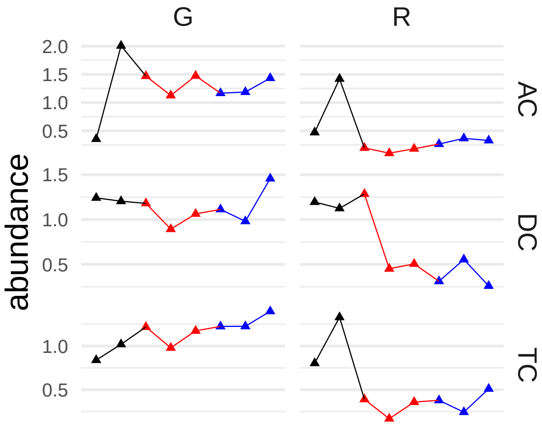


#### [1] "N-stearoyltaurine"

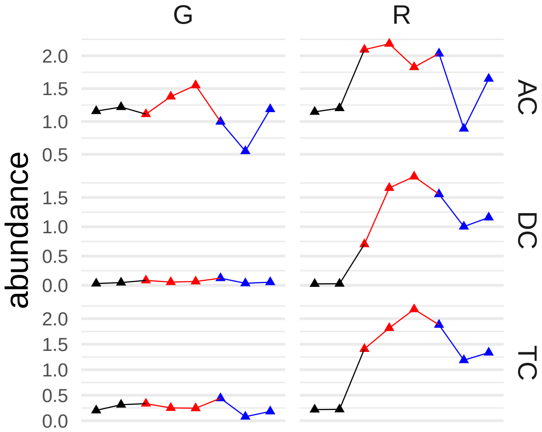


#### [1] "glucuronate"

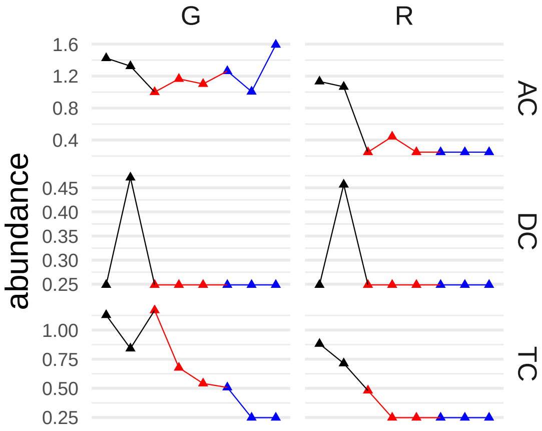


#### [1] "3-hydroxypyridine"

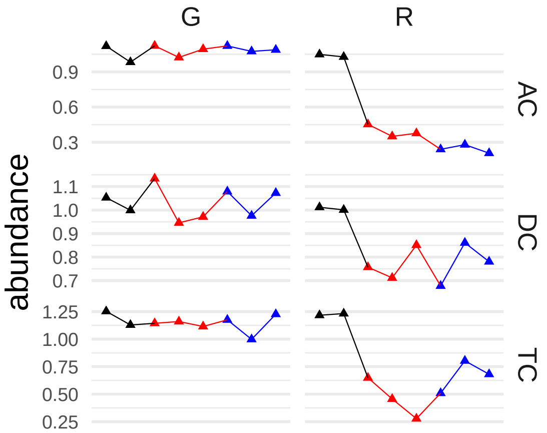


#### [1] "6-phosphogluconate"

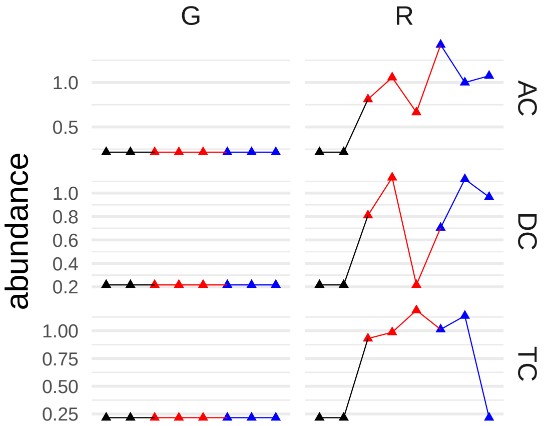


#### [1] "adenosine 5'-monophosphate (AMP)"

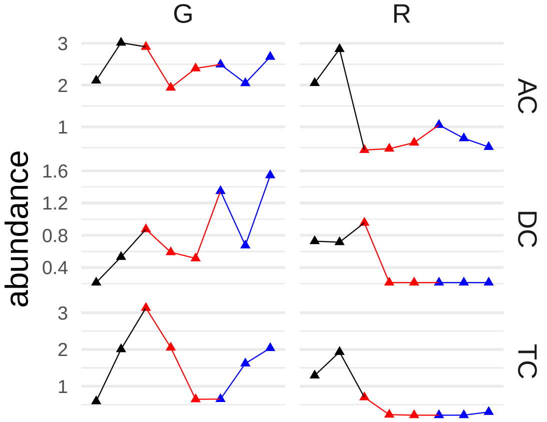


#### [1] "serine"

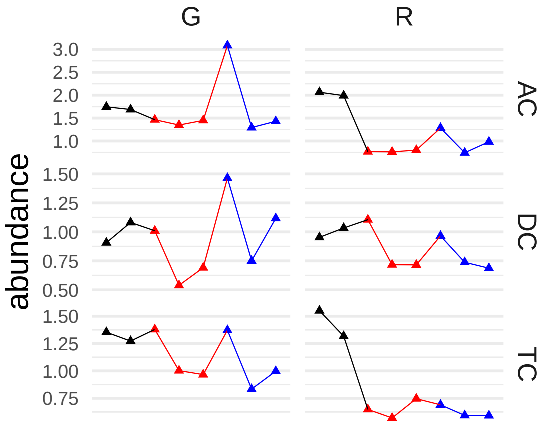


```
## [1] "1-stearoyl-2-oleoyl-GPC (18:0/18:1)"
```

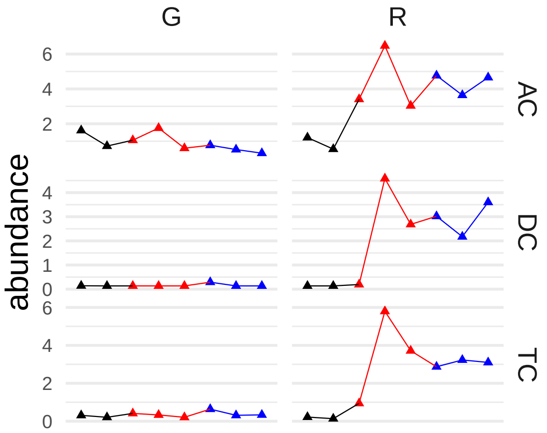
